# CD44 cross-linking promotes *Plasmodium falciparum* invasion

**DOI:** 10.1101/2025.07.30.667750

**Authors:** Angel K. Kongsomboonvech, Stephen W. Scally, Yann Le Guen, Praveesh Valissery, Nichole D. Salinas, Alan F. Cowman, Niraj H. Tolia, Elizabeth S. Egan

**Affiliations:** Department of Pediatrics, Stanford University School of Medicine, Stanford, CA; The Walter and Eliza Hall Institute of Medical Research, Parkville, Australia; Quantitative Sciences Unit, Department of Medicine, Stanford University School of Medicine, Stanford, CA, USA; Host-Pathogen Interactions and Structural Vaccinology Section, Laboratory of Malaria Immunology and Vaccinology, National Institute of Allergy and Infectious Diseases, National Institutes of Health, Bethesda, MD; Department of Microbiology & Immunology, Stanford University School of Medicine, Stanford, CA; Chan Zuckerberg Biohub-San Francisco, San Francisco, CA

## Abstract

The ability of the malaria parasite *Plasmodium falciparum* to invade and replicate asexually within human red blood cells (RBCs) is central to its pathogenicity, accounting for hundreds of thousands of deaths each year. RBC invasion is a multi-step process involving several host-parasite interactions, yet the host factors acting during invasion remain underexplored, largely due to the intractability of mature enucleated RBCs. The transmembrane protein CD44 was identified as a host factor for *P. falciparum* invasion through a forward genetic screen using genetically modified RBCs derived from primary human hematopoietic stem cells. Here, we identify an anti-CD44 monoclonal antibody, BRIC 222, that significantly promotes *P. falciparum* invasion, and demonstrate that its effect is mediated through CD44 cross-linking. CD44 cross-linking induced changes in the phosphorylation of RBC cytoskeletal proteins, consistent with a proposed role for CD44 as a co-receptor during invasion. CD44 cross-linking also altered the RBC membrane, increasing the accessibility of several surface proteins, including the essential invasion receptor Basigin. The parasite ligand Erythrocyte Binding Antigen-175 (EBA-175), which interacts with CD44, enhanced *P. falciparum* invasion and induced RBC membrane changes similarly to BRIC 222. Moreover, both BRIC 222 and EBA-175 increased binding of the PfRH5/PCRCR invasion complex to Basigin, an interaction known to be essential for invasion. We propose that CD44 cross-linking, potentially by EBA-175, serves to coordinate and enhance downstream ligand-receptor interactions and to promote signaling to the host cell cytoskeleton, making RBCs more permissive to *P. falciparum* invasion.

## INTRODUCTION

Malaria remains a global threat with an estimated 249 million cases and 608,000 deaths per year^1^. The disease is caused by *Plasmodium* spp. parasites, with most cases of severe, multi-system disease attributable to *Plasmodium falciparum*. Clinical manifestations occur when *P. falciparum* invades and replicates asexually in red blood cells (RBCs). Every 48 hours, a single intracellular parasite can generate up to 32 daughter merozoites, which then egress and re-invade new RBCs, driving exponential infection^2,3^.

*P. falciparum* invasion is a rapid, multi-step process involving several parasite ligands and host receptors. Parasite surface ligands from the erythrocyte-binding like (EBL) and the *P. falciparum* reticulocyte-binding-like protein homolog (PfRH) families localize at the apical end of the invasive merozoite, where their interactions with host receptors mediate adhesion, apical reorientation, and RBC membrane deformation^2–4^. The only host-parasite interaction shown to be uniquely essential for invasion involves the pentameric *P. falciparum* protein complex PCRCR, which binds to RBC Basigin *via* the parasite ligand PfRH5^5–9^. The binding of PCRCR to Basigin is associated with a calcium spike and is thought to trigger rhoptry organelle secretion and assembly of the moving junction, a critical structure for subsequent parasite-powered internalization into the host RBC^10–12^.

Recently, a forward genetic shRNA screen using cultured red blood cells (cRBCs) derived *ex vivo* from primary human hematopoietic stem cells (HSPCs) identified the transmembrane protein CD44 as a host factor for *P. falciparum*^13^. A role for CD44 in invasion was confirmed through *P. falciparum* invasion assays, where both CD44-depleted and CD44-null cRBCs—generated by CRISPR/Cas9—underwent normal differentiation and enucleation but exhibited reduced susceptibility to *P. falciparum* invasion as compared to isogenic wildtype cRBCs^13,14^. Two established *P. falciparum* invasion ligands from the EBL family, EBA-175 and EBA-140, which bind to the receptors glycophorin A and glycophorin C, respectively^15–17^, were also found to interact directly with CD44^14^. Recombinant EBA-175 clusters uninfected RBCs, enhancing parasite growth and protecting against diverse neutralizing antibodies^18^. Several studies have reported that the interaction of EBA-175 with RBCs alters host cell deformability and modulates phosphorylation of RBC cytoskeletal proteins, suggesting a role for signaling during invasion^19,20^. Phosphoprotein profiling indicated that CD44 likely acts as a co-receptor with the glycophorins to transmit EBA-induced signals to the RBC cytoskeleton, which may facilitate invasion^14^.

Here, we investigated CD44 as a potential target for invasion inhibition using a panel of anti-CD44 monoclonal antibodies. We discovered that none of the antibodies blocked invasion, yet one antibody— BRIC 222—specifically promoted *P. falciparum* invasion through CD44 cross-linking. Both CD44 cross-linking and EBA-175 altered the accessibility of multiple RBC surface proteins, including the essential receptor Basigin, and modulated the phosphorylation of RBC cytoskeletal proteins. Together, these findings support a model in which CD44 cross-linking coordinates two steps of invasion that have been previously considered independent: merozoite apical reorientation and discharge of the rhoptry organelles.

## MATERIALS AND METHODS

### *P. falciparum* culture

*Plasmodium falciparum* strains 3D7, W2mef, D10, and 7G8 were cultured in de-identified human RBCs (Stanford Blood Center) at 2% hematocrit (hct), in complete RPMI media (RPMI-1640 with 25 mM HEPES, 50 mg/L hypoxanthine, 2.42 mM sodium bicarbonate, and 0.5% Albumax) at 37°C in 5% CO_2_ and 1% O_2_ gas mixture.

### *P. falciparum* invasion assays

RBCs from de-identified donors (Stanford Blood Center) were infected with magnet-purified schizont-stage parasites at starting parasitemia ∼1% and final 0.5% hematocrit. Antibodies used included: BRIC 170, BRIC 222, BRIC 222 F(ab), BRIC 235, KZ1, KZ1 F(ab) (all from IBGRL) and IM7 (BioLegend). All antibodies were first dialyzed into incomplete RPMI before use in cellular assays. RII EBA-175 was generated as previously described^21^. Assays were performed in three technical replicates at a volume of 100 µl per well and incubated at 37°C in 5% CO_2_ and 1% O_2_. After overnight incubation, the ring-stage parasitemia was measured by flow cytometry after staining with SYBR green I nucleic acid at 1:2000 dilution (Invitrogen). Data analysis was performed with FlowJo v.10.8.1.

### RBC agglutination (clustering) assays

2×10^6^ RBCs from de-identified donors were incubated with either 25 μg/ml of indicated dialyzed antibodies or 2 μM RII EBA-175 in 50 μl final volume while on rotation for one hour or 30 minutes respectively, in RPMI media supplemented with 10% heat-inactivated fetal bovine serum (HI-FBS) and 100 mM NaCl. After washing, agglutination was quantified using an LSR II flow cytometer (Becton Dickinson) by gating for size (FSC) and granularity (SSC). Analysis of the flow cytometry data was performed using FlowJo v.10.8.1. Measurements of RBC cluster events was confirmed by live cell imaging using a Keyence BZ-X fluorescence microscope.

### Flow cytometry-based RBC membrane profiling

RBCs from de-identified donors were incubated with 25 μg/ml of dialyzed BRIC 222 or BRIC 222 F(ab) for one hour or with 2 μM RII EBA-175 for 30 minutes, in RPMI supplemented with 10% HI-FBS and 100 mM NaCl at 0.4% hct. After washing, RBCs were incubated with primary antibodies for one hour at room temperature. Antibodies used were: anti-BSG-APC mouse monoclonal (clone TRA-1-85, R&D Systems), 1:10; anti-BSG rabbit polyclonal (Proteintech, #11989-1-AP), 1:100; CD235a-FITC (clone 2B7, STEMCELL Technologies), 1:50; BRIC 10-FITC (Santa Cruz Biotechnology, #sc-59183), 1:20,000; CD47-PE (clone CC2C6, BioLegend), 1:20; SLC14A1-APC (clone 888418, R&D Systems), 1:8, and CD151-APC (clone 50-6, Invitrogen), 1:10. Where indicated, samples were incubated for one hour with secondary antibody goat anti-rabbit Alexa Fluor 647 after washing (Invitrogen, #A21244), 1:1000. Phosphatidylserine (PS) externalization was measured using Annexin V-APC (Invitrogen, #R37176), following the manufacturer’s protocol. Flow cytometry was performed with a MACSQuant (Miltenyi Biotec) and analyzed with FlowJo v.10.8.1.

### PCRCR binding assay

The PCRCR binding assay was performed essentially as previously described^8^. The PCRCR complex was assembled with equimolar 400 nM of PfPTRAMP-CSS, PfRIPR, PfCyRPA, and PfRH5 proteins by incubating all proteins together in 1% BSA/PBS at room temperature for an hour. 1×10^7^ RBCs from de-identified donors were stimulated with 25 μg/ml BRIC 222 in 1% BSA/PBS at 1% hct in 100 μl for one hour or with 1.5 μM RII EBA-175 in RPMI media with 10% HI-FBS and 100 mM NaCl at 1% hct in 100 μl for 30 minutes, both at room temperature. After washing, the cells were then incubated with 400 nM pre-assembled PCRCR complex, for one hour at room temperature. To detect binding, the samples were incubated with anti-PfRH5 human antibody clone R5.011^22^ (generated at WEHI from published sequence) at 10 μg/ml for one hour at room temperature, followed by goat anti-Human IgG (H+L) Superclonal™ Secondary Antibody, Alexa Fluor™ Plus 488 (Invitrogen, #A56021) at 1:1000 dilution for one hour at room temperature in the dark. PCRCR binding was quantified by flow cytometry. Data analysis was performed with FlowJo v.10.8.

### PfRH5 binding assay

1×10^7^ RBCs from de-identified donors were stimulated with 2 μM RII EBA-175 in RPMI media with 10% HI-FBS and 100 mM NaCl in 100 μl volume at 1% hct for 30 minutes at room temperature. After washing, the cells were incubated with 400 nM PfRH5 for 1 hour at room temperature. To detect binding, cells were incubated with anti-PfRH5 antibody (5A9-488^23^), 0.2 mg/ml, for one hour at room temperature, and analyzed by flow cytometry. Data analysis was performed with FlowJo v.10.8.1.

### Immunofluorescence assays

Immunofluorescence assays were performed as previously described, with some modifications^24^. RBCs were fixed with 4% paraformaldehyde and 0.0075% glutaraldehyde in PBS for 20 minutes, blocked in 3% BSA/PBS, and then incubated overnight at 4°C with 1:1000 anti-CD44 antibody (BRIC 235, IBGRL) in 3% BSA/PBS. After washing, cells were incubated in 1:2000 Goat anti-Mouse IgG (H+L) Cross-Adsorbed Secondary Antibody, Alexa Fluor™ 488 (Invitrogen) for one hour at room temperature. After washing, the cells were stained with 1:10 anti-BSG-APC (clone TRA-1-85, R&D systems) in 3% BSA/PBS. Images were acquired using a 60X objective on an abberior confocal microscope. Image processing was performed using FiJi with custom scripts, and colocalization analysis was conducted using JaCoP^25^.

### Phosphoproteome-profiling with 2D-DIGE

1×10^9^ RBCs from de-identified donors were pre-treated with phosphatase inhibitors (Sigma Phosphatase Inhibitor Cocktail 2 & 3) at 50% hct, 200 μl final volume in Krebs-Ringer buffer, at 37°C for 30 minutes. Samples were then stimulated with 250 μg/ml BRIC 222 or BRIC 222 F(ab) in Krebs-Ringer buffer at 37°C for one hour. Ghost RBCs were generated from these samples by lysis in ice-cold Complete Lysis Buffer (5 mM sodium phosphate buffer at pH 7.5, 1 mM EDTA, Halt protease/phosphatase inhibitor) at 1:35 sample-to-buffer volumes for 10 minutes followed by extensive washing. The ghost samples were processed at Applied Biomics, Inc (Hayward, CA) for two-dimensional difference gel electrophoresis (2D-DIGE) and DeCyder analysis (DeCyder 2D software version 6.5 from GE Healthcare). Spots with highest difference in phosphorylation between BRIC 222 and BRIC 222 F(ab) stimulation were picked and the associated proteins were identified through mass spectrometry by Applied Biomics, Inc.

### Statistical Analysis

All statistical analyses were performed with GraphPad Prism version 10. Statistical tests included one-way ANOVA, Friedman test, Kruskal-Wallis, Wilcoxon matched-pairs signed rank test and paired t-test, and are indicated for each experiment presented. The two-stage two-step method of Benjamini, Kreiger, and Yekutieli was used for false discovery rate (FDR) correction.

## RESULTS

### CD44 cross-linking promotes *Plasmodium falciparum* invasion

We previously reported that CD44-null cultured red blood cells (cRBCs) derived *ex vivo* from primary human hematopoietic stem/progenitor cells (HSPCs) are less susceptible to *P. falciparum* invasion as compared to isogenic wildtype (WT) RBCs^13,14,26^. Since CD44 is a cell surface protein with a large extracellular domain, we sought to determine whether blocking CD44 would impede *P. falciparum* invasion. To test this, we selected four anti-CD44 monoclonal antibodies (mAbs) targeting distinct epitopes within the extracellular domain of CD44: BRIC 222, BRIC 235, KZ1, and IM7 (Fig. 1A), and tested them in parasite invasion assays using *P. falciparum* strain 3D7. IM7 has been shown to block the interaction between CD44 and hyaluronic acid in other cell types^27,28^. BRIC 170 binds to the cytoplasmic domain of the protein Band 3 and served as a non-targeting control antibody. Late-stage *P. falciparum* schizonts were incubated overnight with acceptor RBCs in the presence of increasing concentrations of each antibody, and the invasion efficiency was measured by flow cytometry after staining with a DNA dye to identify parasitized cells, and confirmed with blood smears. Unexpectedly, none of the antibodies impaired *P. falciparum* invasion. Instead, we observed that the BRIC 222 mAb significantly promoted invasion resulting in up to a ∼80% increase in a dose-dependent manner (Fig. 1B). IM7 produced a more modest enhancement (up to ∼20% increase) while the other anti-CD44 mAbs, BRIC 235 and KZ1, had no detectable effect on parasite invasion (Fig. 1B).

**Figure 1.**
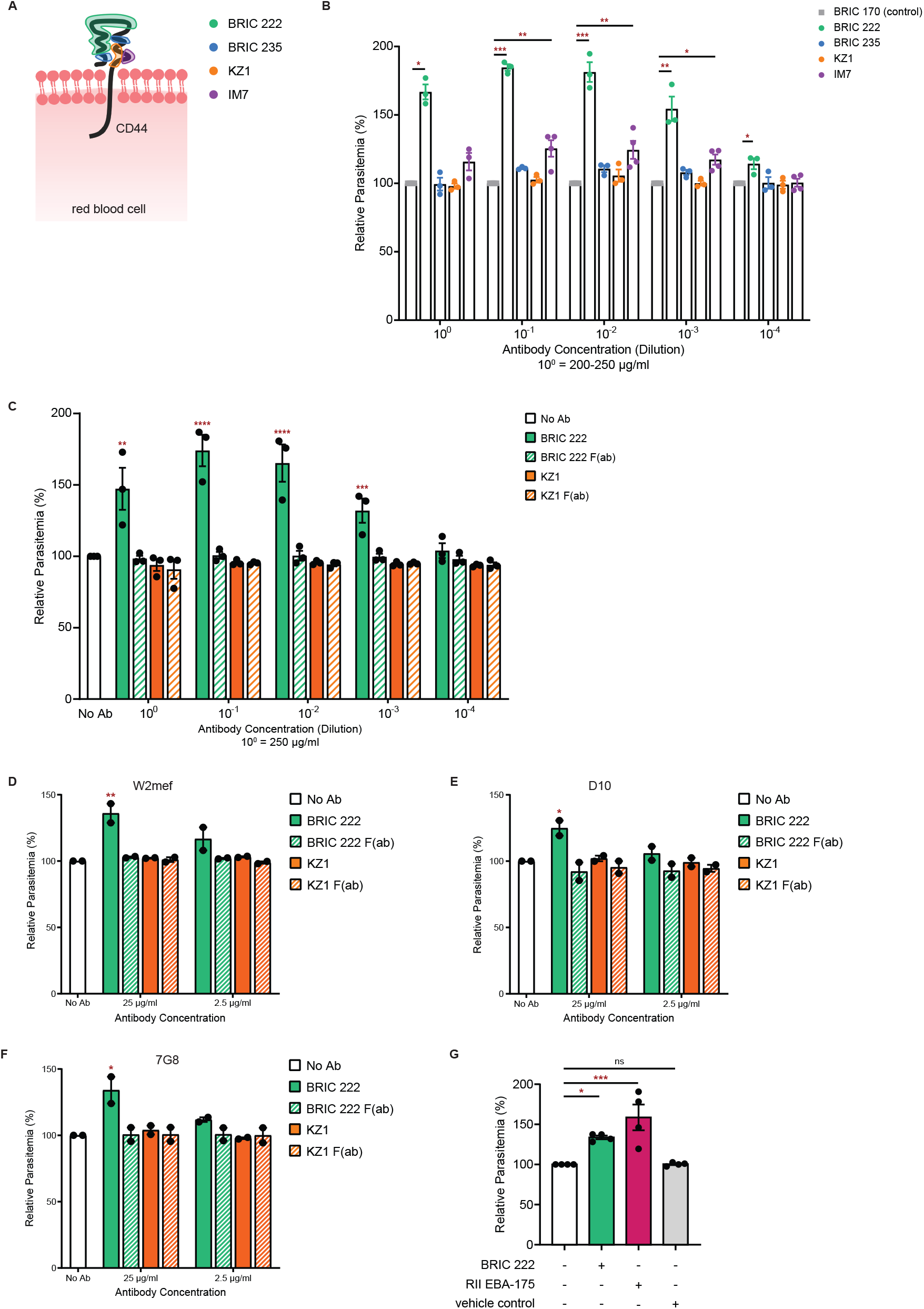
Cross-linking of RBC CD44 promotes *Plasmodium falciparum* invasion. **(A)** Approximate target epitope sites for each anti-CD44 antibodies (BRIC 222, BRIC 235, KZ1, IM7), highlighted and color coded. **(B)** Invasion efficiency of *Plasmodium falciparum* strain 3D7 in presence of anti-CD44 monoclonal antibodies relative to an isotype control (BRIC 170). The parasitemia was measured ∼21 hours post-infection, and normalized to isotype control. Each dot represents one biological replicate (N = 3-4 biological replicates (individual donors); n = 3 technical replicates). Error bars indicate SEM. Statistical analysis: Kruskal-Wallis with FDR correction; * p ≤ 0.05, ** p ≤ 0.01, *** p ≤ 0.001. **(C-F)** Invasion efficiency of *P. falciparum* strains 3D7 (C), W2mef (D), D10 (E), and 7G8 (F) in the presence of the indicated antibodies, relative to no antibody. Antibody concentration at 10^0^ is 25 µg/ml for all except the BRIC 222 F(ab) used in W2mef assay (34 µg/ml). The parasitemia was measured ∼21 hours post-infection, and normalized to an infection with no antibody (No Ab). Each dot represents one biological replicate (N = 2-3 biological replicates; n = 3 technical replicates). Error bars indicate SEM. Statistical analysis: One-way ANOVA with FDR correction; * p ≤ 0.05, ** p ≤ 0.01, *** p ≤ 0.001, **** p ≤ 0.0001. **(G)** Invasion efficiency of *P. falciparum* strain 3D7 in presence of 0.75 µM RII EBA-175, 25 µg/ml BRIC 222, or vehicle control, relative to no additive. Each dot represents one biological replicate (N = 4 biological replicates; n = 3 technical replicates). Error bars represent SEM. Statistical analysis: One-way ANOVA with FDR correction; * p ≤ 0.05, *** p ≤ 0.001, ns: non-significant.

Since IgG molecules can mediate cross-linking of their target antigens, we hypothesized that the increased parasite invasion observed with BRIC 222 was due to CD44 cross-linking. To test this, we compared the parasite invasion efficiency in the presence of BRIC 222 *versus* its antigen-binding fragment BRIC 222 F(ab) that can bind CD44 but not cross-link it. We observed that BRIC 222 F(ab) had no effect on parasite invasion even at the highest concentrations tested (Fig. 1C). This indicates that the enhanced parasite invasion observed with BRIC 222 is dependent on CD44 cross-linking, rather than solely on epitope binding.

To determine if the increased invasion phenotype observed with BRIC 222 was strain-specific, we tested three other *P. falciparum* strains that originate from different geographic areas and employ distinct invasion pathways: W2mef, D10, and 7G8^29–31^. We found that incubation with BRIC 222, but not BRIC 222 F(ab), enhanced the invasion of each of the tested *P. falciparum* strains, suggesting that CD44 cross-linking promotes invasion in a strain-transcendent manner (Fig. 1D-F). This is consistent with prior work using an erythroid cell line implicating CD44 in a strain-transcendent role in *P. falciparum* invasion^26^.

### EBA-175 enhances invasion similarly to BRIC 222

The *P. falciparum* invasion ligand EBA-175 has been shown to interact with CD44 *in vitro*^14^ and recombinant EBA-175 (rEBA-175) has also been demonstrated to promote parasite growth over several cycles^18^. Our finding that BRIC 222 promotes *P. falciparum* invasion through cross-linking CD44 raised the hypothesis that *P. falciparum* EBA-175 may also specifically promote invasion. To test whether EBA-175 can enhance invasion, we performed *P. falciparum* invasion assays with recombinant EBA-175 RII region (RII EBA-175), the region that interacts with RBCs^21^ and is known to directly interact with CD44^14^. We found that incubation with RII EBA-175 led to ∼50% increase in *P. falciparum* invasion relative to the control (Fig. 1G). These results collectively show that both EBA-175 and BRIC 222 enhance *P. falciparum* invasion.

### RBC agglutination induced by CD44 cross-linking is not directly correlated with invasion

Prior work by Paing *et al*. demonstrated that rEBA-175 induces clustering (agglutination) of uninfected RBCs^18^. This observation was proposed as a mechanism for the underlying ability of rEBA-175 to enhance parasite growth *in vitro*; by clustering RBCs together in close proximity, rEBA-175 may facilitate merozoite reinvasion^18^. Notably, BRIC 222 is reported to be a hemagglutinin^32^, raising the possibility that it may enhance invasion *via* RBC agglutination. We used flow cytometry to quantify RBC agglutination induced by RII EBA-175, BRIC 222, or the other anti-CD44 antibodies (Fig. 2A-B). We found that BRIC 222 induced RBC agglutination, or cluster events, in ∼7% of the population, similar to rEBA-175 (Fig. 2A-C). In contrast, IM7 induced a milder agglutination phenotype (∼2-3% of the population), whereas cells incubated with BRIC 235 and KZ1 did not show detectable agglutination, resembling unstimulated cells (Fig. 2A-C). To determine if the agglutination phenotype observed with BRIC 222 was due to CD44 cross-linking, we also tested BRIC 222 F(ab). The results showed that BRIC 222 F(ab) does not induce agglutination, suggesting that the phenotype is indeed due to CD44 cross-linking (Fig. 2D).

**Figure 2.**
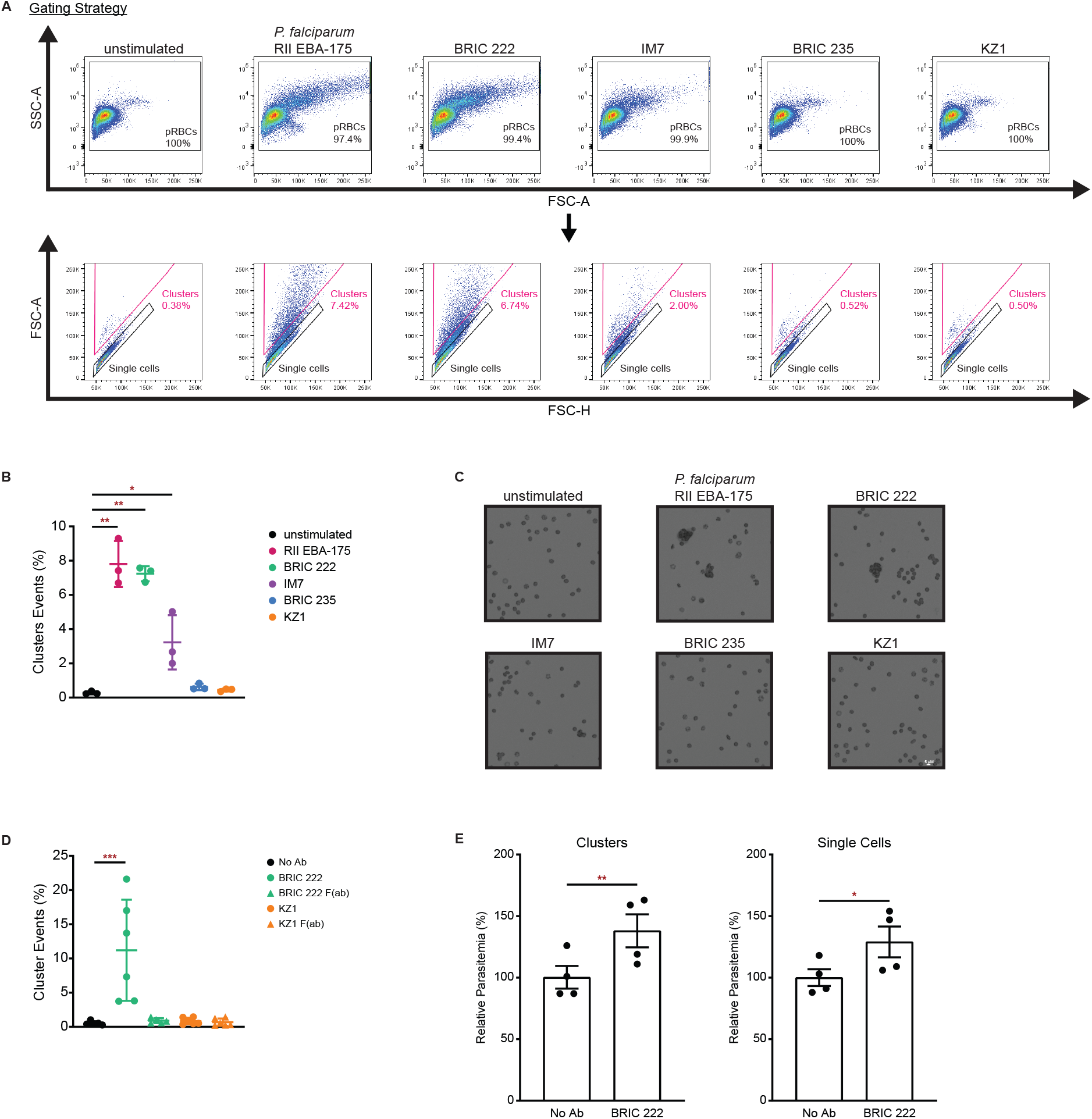
RBC agglutination is not directly correlated with enhanced *P. falciparum* invasion. **(A)** Gating strategy for flow cytometry analysis of RBC clusters events from a representative experiment. RBC agglutination/clustering of three cells or more is denoted as “Clusters” in orange. **(B)** RBCs were treated with 25 µg/ml anti-CD44 monoclonal antibodies or 2 µM RII EBA-175 and assessed for agglutination/clustering by flow cytometry, measured by forward scatter (size). Frequency of cluster events from three donors with average ± SD is shown. Statistical analysis: Kruskal-Wallis with FDR correction; * p ≤ 0.05, ** p ≤ 0.01. **(C)** Representative images of RBCs treated with 25 µg/ml anti-CD44 monoclonal antibodies (BRIC 222, IM7, BRIC 235, KZ1) or 2 µM RII EBA-175, taken with 20X objective. **(D)** RBCs were treated with 25 µg/ml of the indicated monoclonal antibodies or their respective F(ab)s. Average frequency of cluster events from 5-6 donors ± SD is shown. Statistical analysis: Kruskal-Wallis with FDR correction; *** p ≤ 0.001. **(E)** The relative invasion efficiency within populations of “Clusters” or “Single cells” was measured by flow cytometry and normalized to the mean of the no antibody (No Ab) condition. Each dot represents an individual biological replicate (N = 4 biological replicates; n = 3 technical replicates). Error bars represent SEM. Statistical analysis: paired t-test, one-tailed; ** p ≤ 0.01, * p ≤ 0.05.

As the agglutination phenotypes associated with the different monoclonal antibodies correlated with their respective invasion phenotypes, we next sought to determine if the enhanced invasion observed with BRIC 222 is attributable to RBC agglutination. We performed *P. falciparum* invasion assays in the presence or absence of BRIC 222 and measured the resulting parasitemia separately in non-agglutinated singlet cells *versus* agglutinated clusters by flow cytometry. We observed that the addition of BRIC 222 led to enhanced *P. falciparum* invasion in both non-agglutinated singlet cells and in the clusters (Figs. 2E and S1). Collectively, these findings illustrate that even though BRIC 222 can induce RBC agglutination, this does not fully explain the phenotype of increased parasite invasion because only a small proportion of cells were agglutinated, and *P. falciparum* invasion was similarly enhanced in the non-agglutinated singlet cells (Figs. 2E and S1).

### CD44 cross-linking alters phosphorylation of RBC cytoskeletal proteins

Incubation of RBCs with rEBA-175 has been shown to induce cellular biophysical changes as well as altered phosphorylation of host cytoskeletal proteins^19,20^, some of which appear to be dependent on CD44^14^. To determine if CD44 cross-linking alters RBC cytoskeletal protein phosphorylation, we performed phosphoproteome-profiling using two-dimensional difference gel electrophoresis (2D-DIGE) on RBCs stimulated with BRIC 222 or BRIC 222 F(ab). Comparison of the phosphorylation status of membrane ghost proteins from BRIC 222 F(ab)- *versus* BRIC 222-stimulated RBCs revealed differentially-phosphorylated spots with ≥1.5 or ≤0.67 fold-change in phosphorylation (Figs. 3A and S2). We selected five spots that were highly differentially phosphorylated between the BRIC 222 F(ab) and BRIC 222 conditions (ratio of 0.2 or lower) for identification by mass spectrometry. This analysis identified several proteins previously implicated in the RBC cytoskeletal remodeling by *Plasmodium* invasion ligands, including ankyrin-1, alpha and beta spectrin, alpha adducin, and Band 3^14,19,20,33^ (Fig. 3B). These findings indicate that CD44 cross-linking alters the phosphorylation status of host RBC cytoskeletal proteins, similar to EBA-175, suggesting that CD44 may influence *P. falciparum* invasion by acting as a sensor of the extracellular environment.

**Figure 3.**
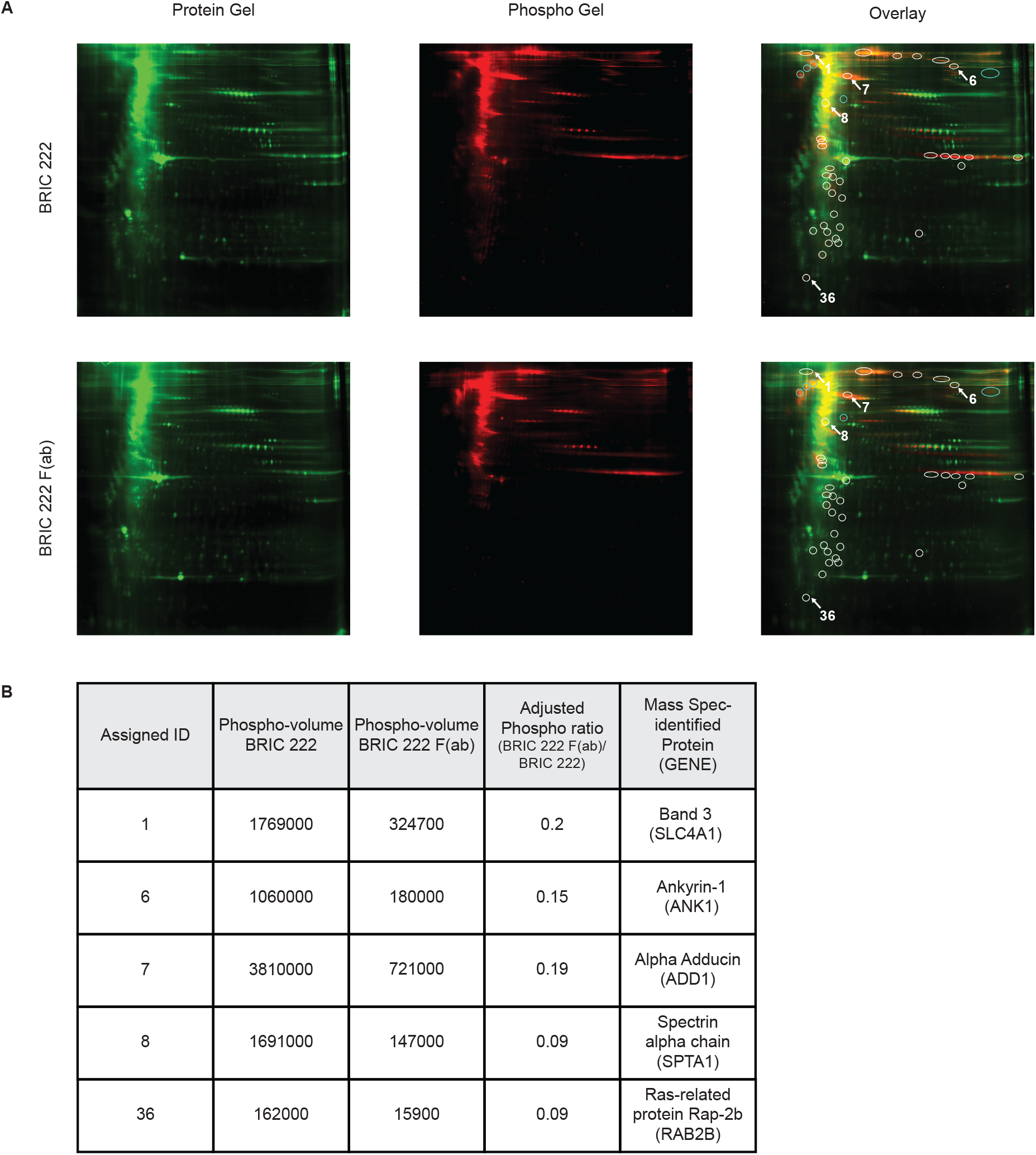
CD44 cross-linking induces phosphorylation of RBC cytoskeletal proteins RBCs were stimulated with BRIC 222 or BRIC 222 F(ab) at 25 µg/ml, processed into ghosts, and used for 2D-DIGE. **(A)** Total Protein and Phosphoprotein gels, along with the overlays of each condition are shown. Spots with differential phosphorylation, with ≥1.5 or ≤0.67 fold-change in phosphorylation, between the BRIC 222 F(ab)- and BRIC 222-stimulated conditions are circled. White circles indicate higher phosphorylation in BRIC 222-stimulated samples; blue circles indicate higher phosphorylation in BRIC 222 F(ab)-stimulated samples. **(B)** Top phosphorylated candidates, numbered in (A) with their respective phospho-volume. Adjusted phospho-ratio indicates phosphorylation ratio adjusted based on protein expression. The candidate proteins were identified by mass spectrometry, with the associated gene indicated in the parentheses.

### CD44 cross-linking alters the RBC plasma membrane

In other cells, CD44 cross-linking has been shown to induce a myriad of changes, such as regulating immunomodulatory functions, modulating signaling pathways, and altering expression or localization of other membrane and cytoskeletal proteins^34–39^. While enucleated RBCs cannot synthesize new proteins, we hypothesized that CD44 cross-linking may modify the architecture of the plasma membrane in a manner that impacts receptor accessibility to invading parasites. To test this idea, we stimulated RBCs with BRIC 222 or BRIC 222 F(ab) and then quantified the relative abundance of specific RBC membrane proteins by flow cytometry. We observed that CD44 cross-linking by BRIC 222 resulted in increased surface detection of all membrane proteins tested, including several known to be important for invasion, such as Basigin (BSG), glycophorin A (GYPA), glycophorin C (GYPC), and others with no known role in invasion, including CD47, SLC14A1 and CD151^13^ (Figs. 4, S3, and S4). We also measured phosphatidylserine (PS) exposure and found that BRIC 222 increased PS externalization while BRIC 222 F(ab) had no effect (Fig. S4C). Collectively, these results indicate that CD44 cross-linking by BRIC 222 induces global changes to the RBC membrane.

**Figure 4.**
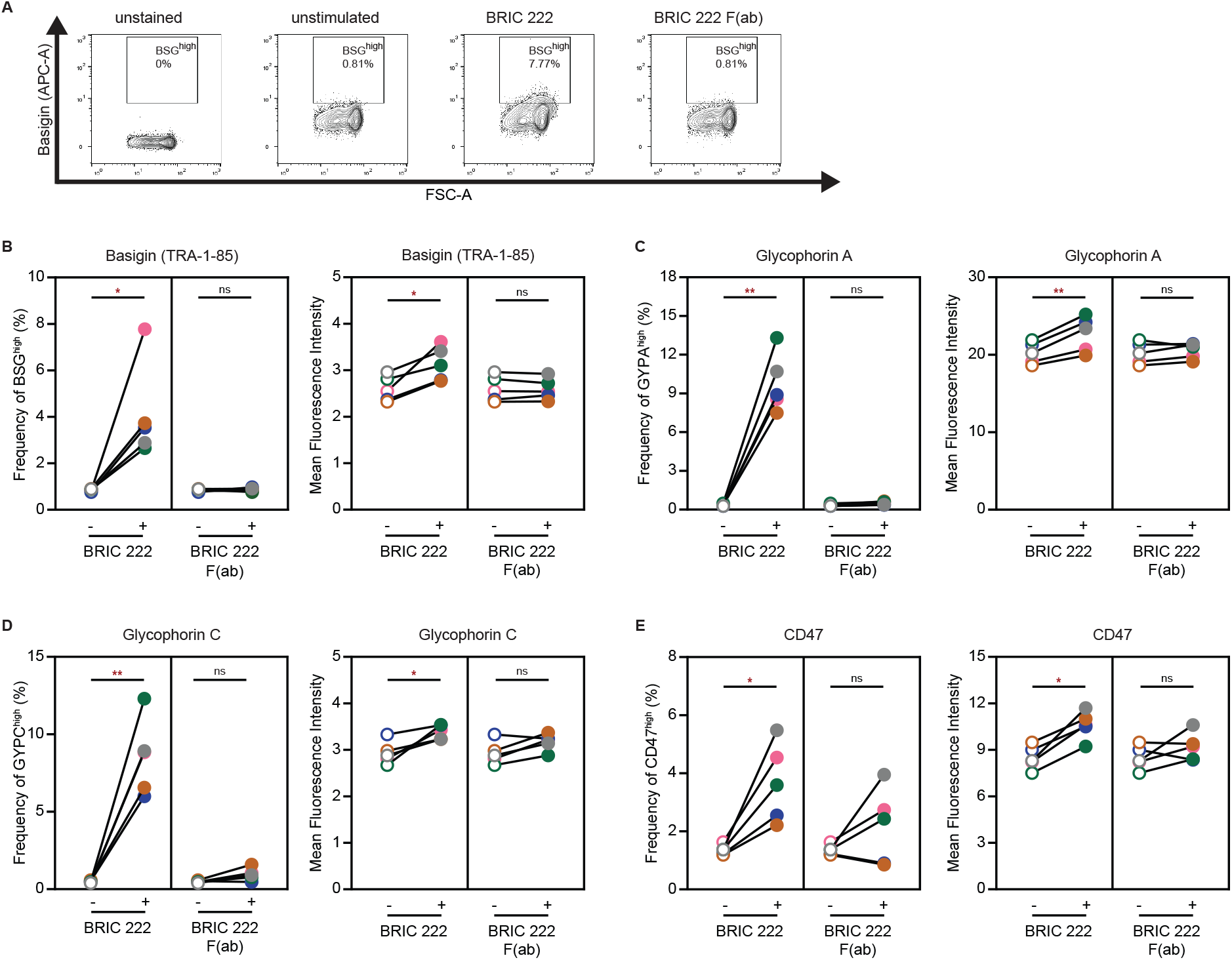
CD44 cross-linking alters the accessibility of Basigin and other RBC membrane proteins RBCs were stimulated with BRIC 222 or BRIC 222 F(ab) at 25 µg/ml and then assessed for surface protein changes. **(A)** Representative gating for RBC surface Basigin (BSG) detected with monoclonal anti-BSG antibody TRA-1-85, indicating BSG^high^ population. **(B-F)** Frequencies of BSG^high^ (B), GYPA^high^ (C), GYPC^high^ (D), and CD47^high^ (E) as well as the respective MFI of five distinct RBC donors are plotted. For each graph, a color dot represents an individual donor. Statistical analysis: Friedman test with FDR correction; * p ≤ 0.05, ** p ≤ 0.01, ns: non-significant.

### CD44 cross-linking enhances the accessibility of a Basigin epitope critical for invasion

As BSG is known to be an essential receptor for *P. falciparum*^5^ and has been shown to co-exist in a complex with CD44 in other cell types^40,41^, we sought to further assess how CD44 cross-linking alters BSG accessibility. In RBCs, the precise spatial organization of CD44 and BSG on the cell surface is not well-defined, but they have been proposed to interact based on genetic and biochemical evidence^26,42^. Using immunofluorescence assays, we observed some co-localization of BSG and CD44 (Fig. 5A), which is consistent with an interaction. To investigate how the enhanced invasion induced by CD44 cross-linking may relate to BSG accessibility, we stimulated RBCs with our panel of anti-CD44 antibodies and used flow cytometry to detect BSG at the surface with either a polyclonal anti-BSG antibody or the anti-BSG monoclonal antibody TRA-1-85, which has been shown to inhibit invasion^5^. We observed that stimulation with either BRIC 222 or IM7 led to increased binding by the polyclonal anti-BSG antibody (Fig. 5B), suggesting an increased accessibility of Basigin to antibody. Moreover, binding by the monoclonal anti-BSG antibody TRA-1-85 was also enhanced after stimulation with BRIC 222, and to a lesser degree with IM7 (Fig. 5C). Together, these results suggest that CD44 cross-linking enhances the surface exposure of Basigin, specifically at an epitope important for invasion, and correlate positively with the results of the invasion assays (Fig. 1B).

**Figure 5.**
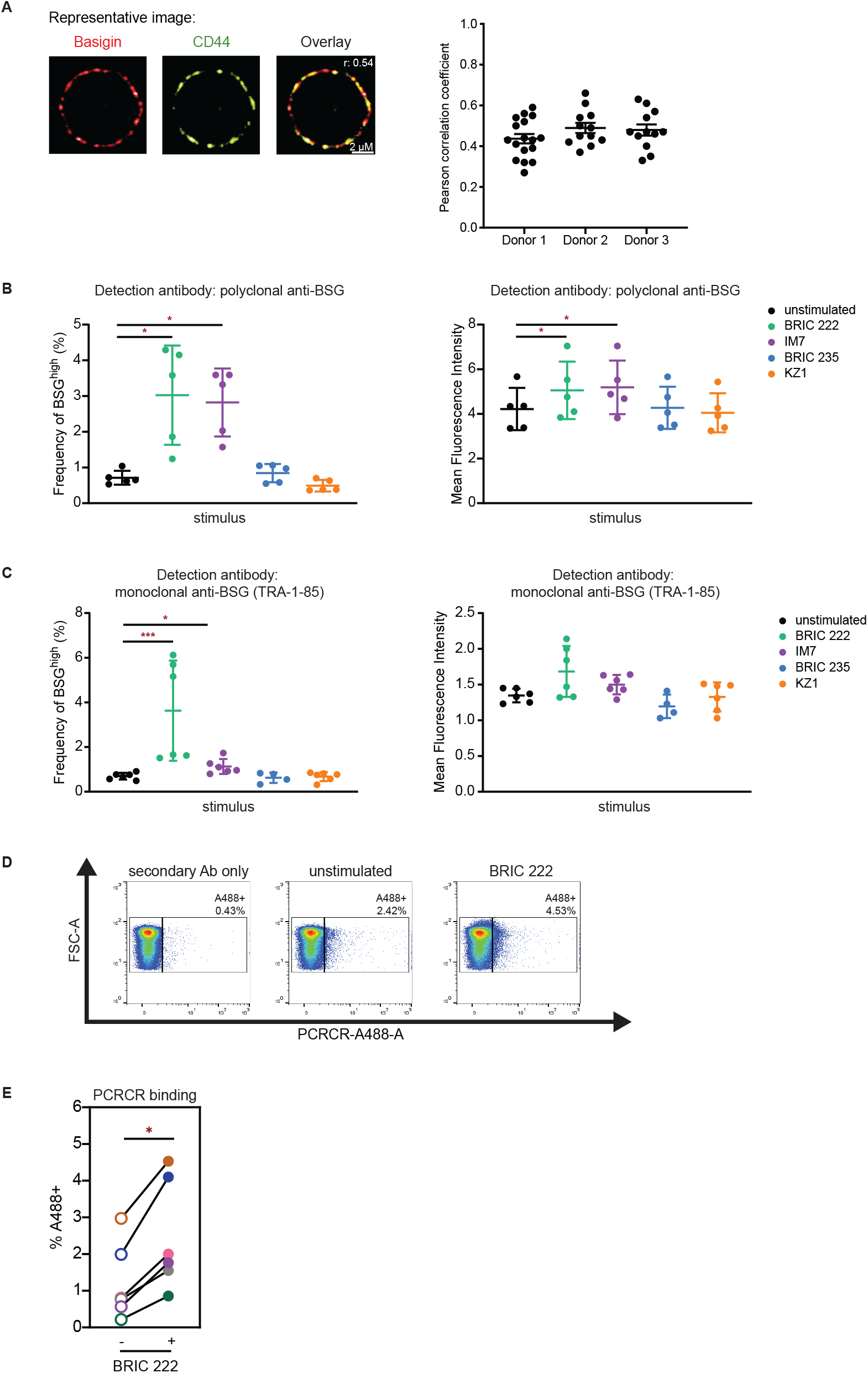
CD44 cross-linking promotes *P. falciparum* PCRCR binding. **(A)** Left: Representative images from immunofluorescence assay for Basigin (BSG) and CD44 on RBCs from an individual donor. Pearson correlation coefficient for co-localization (r) is indicated. Right: Pearson correlation coefficients for co-localization of BSG and CD44 on individual cells from three distinct donors are plotted; each dot represents a single RBC; error bars indicate SEM. **(B)** RBCs were stimulated with the indicated anti-CD44 antibodies (25 µg/ml) and then surface BSG was detected by flow cytometry with anti-BSG polyclonal antibody. Each dot represents an individual donor, and frequency of BSG^high^ (left) and MFI (right) are plotted. The average BSG^high^ or MFI ± SD for five donors is indicated. Statistical analysis: Friedman test with FDR correction; * p ≤ 0.05. **(C)** RBCs were stimulated with the indicated anti-CD44 antibodies (250 µg/ml) and then detected through surface staining of anti-BSG monoclonal antibody TRA-1-85. Frequency of BSG^high^ population (left) and MFI (right) are plotted. Average of 4-6 donors ± SD is indicated; each dot represents one donor. Statistical analysis: Kruskal-Wallis with FDR correction; * p ≤ 0.05, *** p ≤ 0.001. **(D)** PCRCR binding to RBCs was assessed following stimulation with 25 µg/ml BRIC 222. PCRCR binding was detected using anti-PfRH5 (R5.011). Flow cytometry gating strategy for a representative donor is shown. Percentage of PCRCR binding is boxed and indicated as A488+. **(E)** PCRCR binding results from six donors, comparing the unstimulated and BRIC 222-stimulated conditions, are plotted. Each color dot represents an individual donor. Statistical analysis: Wilcoxon matched-pairs signed rank test, one-tailed; * p ≤ 0.05.

Stimulation with BRIC 222 uniquely enhances *P. falciparum* invasion relative to the other anti-CD44 antibodies, and TRA-1-85 is known to bind to an epitope required for invasion. These data suggest that CD44 cross-linking by BRIC 222 may promote invasion by exposing a specific epitope of BSG that enhances the interaction of the merozoite with the RBC. During invasion, the merozoite interacts with BSG *via* its pentameric PCRCR complex, consisting of PfPTRAMP-CSS, PfRIPR, PfCyRPA, and PfRH5^6,8^. To directly test the impact of CD44 cross-linking on PCRCR binding to the RBC, we employed a cellular assay where RBCs were stimulated with BRIC 222 to cross-link CD44, incubated with PCRCR complex, and then PCRCR binding was quantified by flow cytometry. In unstimulated cells, PCRCR binding was detected at a low level, consistent with previous reports^6,8^. After stimulation with BRIC 222, there was an average increase of 2.5-fold in PCRCR binding to RBCs (range 1.53-fold to 4-fold), demonstrating that CD44 cross-linking not only enhances the accessibility of BSG to antibody, but also increases its interaction with PCRCR (Fig. 5D-E, S5). Together, these findings strongly suggest that the enhanced invasion observed for RBCs stimulated with BRIC 222 is due to increased interaction between PCRCR and its receptor BSG on the RBC, at least in part.

As EBA-175 is known to interact with CD44 in addition to its canonical receptor GYPA and can promote *P. falciparum* invasion similarly to BRIC 222, we hypothesized that EBA-175 may cross-link CD44 during parasite infection. To investigate this possibility, we stimulated RBCs with RII EBA-175 and then measured the surface accessibility of the previously mentioned membrane proteins—BSG, GYPA, GYPC, CD47, SLC14A1, and CD151—as well as PS externalization. In addition to the increase in PS externalization, all proteins became more accessible following RII EBA-175 stimulation (Fig. 6A-D; Fig. S6). This suggests that *P. falciparum* RII EBA-175 induces global changes to the RBC membrane, including more exposure of the invasion-specific epitope on BSG, similar to BRIC 222.

**Figure 6.**
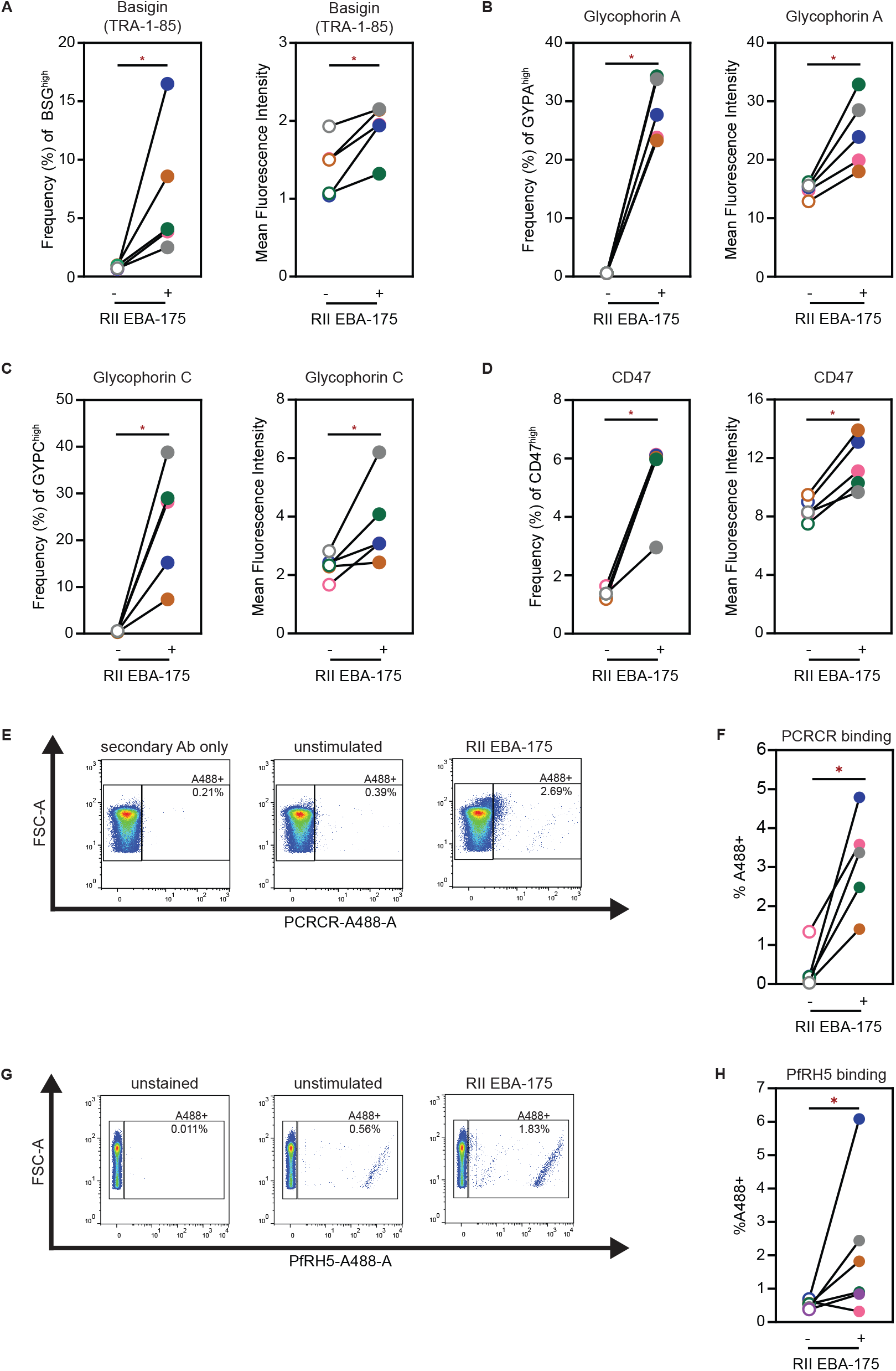
CD44 cross-linking by *P. falciparum* EBA-175 increases PCRCR binding to RBCs. **(A-D)** RBCs were stimulated with 2 µM RII EBA-175. (A) Basigin (BSG) was detected with the anti-BSG monoclonal antibody TRA-1-85. Frequency of BSG^high^ population (left) and MFI (right) of five donors are shown. Frequencies of GYPA^high^ (B), GYPC^high^ (C), CD47^high^ (D) and the respective MFI of five donors are plotted. Each color dot represents an individual donor. Statistical analysis: Wilcoxon matched-pairs signed rank test, one-tailed; * p ≤ 0.05. **(E)** PCRCR binding of RBCs was assessed following 1.5 µM RII EBA-175 stimulation. Representative gating for one donor is shown. Percentage of PCRCR binding is boxed and indicated as A488+. **(F)** PCRCR binding of five donors, comparing unstimulated and EBA-175-stimulated conditions, is plotted. Each color dot represents an individual donor. Wilcoxon matched-pairs signed rank test, one-tailed; * p ≤ 0.05. **(G)** PfRH5 binding on RBCs was assessed following stimulation with 2 µM RII EBA-175. PfRH5 was detected using an anti-PfRH5 antibody conjugated with A488 fluorophore (5A9-488). Representative gating for one donor is shown, with percentage of PfRH5 binding boxed and indicated as A488+. **(H)** Percentage of PfRH5 binding compared to unstimulated condition, across six donors, is plotted. Each color dot represents an individual donor. Statistical analysis: Wilcoxon matched-pairs signed rank test, one-tailed; * p ≤ 0.05.

We next quantified binding of PCRCR to EBA-175-stimulated RBCs. PCRCR binding increased by an average of 35-fold after stimulation with EBA-175 (range 2.7-fold to 112-fold) (Figs. 6E-F, S7). Moreover, RBCs stimulated with EBA-175 also displayed enhanced binding of PfRH5 alone, in absence of the PCRCR complex, with an average increase in binding of 3.7-fold (range 0.5-fold to 8.8-fold) (Fig. 6G-H). Altogether, our data show that both BRIC 222 and EBA-175 lead to more exposure of the invasion-specific epitope of BSG, resulting in enhanced PCRCR/PfRH5 binding to RBCs. Thus, we propose a model in which *P. falciparum* EBA-175 cross-links RBC CD44 upon RBC binding, leading to enhanced interaction between PCRCR and BSG, as well as changes to the RBC cytoskeleton, thereby promoting *P. falciparum* invasion.

## DISCUSSION

During red blood cell (RBC) invasion, *P. falciparum* merozoites progress through a complex series of steps beginning with initial contact and apical reorientation, and culminating in internalization and formation of the parasitophorous vacuole. Facilitated by distinct protein-protein interactions, the steps of invasion have been characterized as independent yet coordinated, but the mechanisms underlying this precise control are unclear. In this study, we investigated the hypothesis that the RBC surface protein CD44 acts as a co-receptor during invasion, by linking the early steps of apical reorientation and RBC deformation (mediated by the interaction of EBA-175 with GYPA) with the subsequent release of the rhoptry contents. Our findings support a model where CD44 cross-linking by the *P. falciparum* ligand EBA-175 primes the RBC membrane to enhance the essential interaction between host BSG and *P. falciparum* PCRCR and promotes signaling to the RBC cytoskeleton, ensuring co-regulation of the sequential steps of invasion.

Here, we initially set out to identify anti-CD44 antibodies capable of blocking *P. falciparum* invasion; however, none of those tested exhibited an inhibitory effect. Instead, one antibody, BRIC 222, strongly enhanced invasion in a strain-transcendent manner. This was unexpected, as to our knowledge all antibodies to RBC receptors that have been previously reported to impact invasion are inhibitory (e.g. anti-BSG, anti-CR1, anti-CD55, anti-GYPA, anti-GYPC)^5,17,43–45^. The phenotype of enhanced invasion suggested that BRIC 222 was engaging and stimulating the ectodomain of CD44, rather than blocking it. BRIC 222 F(ab) had no impact on invasion, indicating that the phenotype observed with BRIC 222 is due to CD44 cross-linking. Together with prior research showing that CD44 is required for efficient invasion of all strains tested^13,14,26^, these data demonstrate that CD44 cross-linking plays a critical and specific role during invasion.

Work by Paing *et al*. suggested that shed EBA-175 promotes *P. falciparum* growth by clustering nearby uninfected RBCs, making them more accessible to invading merozoites^18^. As BRIC 222 is considered a hemagglutinin, we initially investigated whether antibody-induced agglutination could explain the enhanced invasion phenotype observed. We confirmed that BRIC 222 can agglutinate RBCs, but at the antibody concentrations associated with enhanced invasion, most cells in the population remained non-agglutinated. When we directly measured parasitemia in agglutinated cells *versus* the individual, non-agglutinated cells, we found that they both had similarly elevated rates of invasion relative to unstimulated RBCs. This indicates that the enhanced *P. falciparum* invasion observed with BRIC 222 cannot be fully explained by RBC agglutination, prompting us to investigate alternative hypotheses.

Prior observations of altered phosphorylation of RBC cytoskeletal proteins and/or increased membrane deformability upon stimulation with *P. falciparum* invasion ligands imply the existence of RBC signaling pathways that may be exploited by the parasite to establish infection^12,14,19,20,33^. Indeed, phosphorylation of membrane and cytoskeletal proteins is considered a mechanism to regulate properties of the RBC membrane in response to environmental triggers^46^. Previously, we found that EBA-induced phosphorylation of RBC cytoskeletal proteins differed between WT and CD44-null cRBCs, suggesting that CD44 may facilitate signaling during invasion^14^. Our results here provide further support for this idea, as CD44 cross-linking by BRIC 222 alters the phosphorylation of several RBC cytoskeletal proteins, such as ankyrin-1, alpha and beta spectrin, alpha adducin, and Band 3. Recent work from Yong *et al*. further illuminates this finding, as they found that signaling induced by adhesin binding involves activation of the Beta-adrenergic receptor (β2AR), which they demonstrated to be in complex with BSG and CD44^42^. Together, these results suggest that that CD44 cross-linking may enhance invasion by facilitating signaling to the RBC cytoskeleton in a manner that involves interactions with neighboring proteins, such as BSG.

Indeed, the essential receptor BSG does not reside in the RBC membrane in isolation. Instead, it localizes to at least two discrete multi-protein complexes, interacting with the monocarboxylate transporter (MCT1) or with PMCA4, the major calcium transporter of human RBCs. Recent work showed that the binding affinity between PfRH5 and BSG was strengthened when BSG was present within one of these complexes rather than by itself, suggesting that its context within the membrane may have functional consequences for invasion^7^. Another recent study used genetically modified reticulocytes derived from an erythroid cell line, BEL-A, to investigate the co-dependence of BSG and MCT1^47^. The surface expression of BSG was found to be dependent on MCT1 and *vice versa*. Intriguingly, ectopic expression of the BSG extracellular domain fused to a GPI anchor could complement the invasion phenotype of BSG-null cells, suggesting that neither the transmembrane nor cytoplasmic domains of BSG are required for its essential function^47^.

Our data for CD44 are not inconsistent with a model where BSG’s function during invasion is exerted solely by its ectodomain. CD44 cross-linking resulted in increased surface detection of the BSG ectodomain, increased PCRCR/PfRH5 binding, and enhanced invasion; since RBCs cannot synthesize new proteins, this suggests that cross-linking CD44 alters BSG at a structural level. Indeed, our data suggest that CD44 cross-linking promotes invasion by changing the BSG ectodomain in a way that enhances its interaction with PfRH5 and the PCRCR complex. A recent study by Day *et al*. employed surface plasmon resonance to examine the determinants of PfCyRPA binding to membrane ghosts from an erythroid cell line, as PfCyRPA is part of the PCRCR complex. They found that CD44 and BSG contributed to the binding of PfCyRPA to the RBC to a similar degree, likely though specific sialyated residues^48^. Moreover, BSG-null reticulocytes derived from the BEL-A cell line have been shown to have higher levels of surface CD44 as compared to isogenic WT cells, further supporting the idea that BSG and CD44 are both functionally and structurally related^47^.

A pivotal study by Weiss *et al*. defined the sequential steps of *P. falciparum* invasion using reagents to block specific ligand-receptor interactions^11^. Blocking the interaction between BSG and PfRH5 using antibodies inhibited echinocytosis and Ca^2+^ influx into the RBC, both of which are considered measures of rhoptry release and perforation of the erythrocyte membrane. In light of subsequent work establishing PfRH5 as part of the PCRCR complex together with PfCyRPA, PfRipr, PfPTRAMP and PfCSS^7,8,49^, these findings established that binding of PCRCR to BSG is a prerequisite for discharge of the rhoptry organelles and perforation of the host cell membrane during invasion. Our results showing that CD44 cross-linking specifically alters the RBC membrane and increases the accessibility of BSG to PCRCR and PfRH5 suggest that CD44 plays a key, upstream role in modulating the interaction between PCRCR and BSG.

The studies here utilized BRIC 222 antibody to cross-link CD44. What ligand(s) could perform this activity during an actual infection? EBA-175 is the most likely candidate, based on several lines of evidence. First, EBA-175 can directly bind to CD44 in addition to its canonical receptor GYPA, as demonstrated by pull-downs from parasite lysate and confirmed by cellular binding assays using recombinant proteins^14^, and functions as a dimer^50^. Second, EBA-175 can enhance *P. falciparum* invasion, mimicking the phenotype observed for BRIC 222-induced cross-linking of CD44. Third, incubation of RBCs with both EBA-175 and BRIC 222 leads to altered phosphorylation of RBC cytoskeletal proteins. Fourth, both EBA-175 and BRIC 222 change the RBC membrane, increasing the accessibility of BSG and the subsequent PCRCR/PfRH5 binding to RBCs, a crucial step for successful invasion. While these data support the idea that EBA-175 serves to cross-link CD44 during invasion, ultimately structural studies will be needed to fully characterize the interactions between EBA-175, CD44, the glycophorins, and BSG. Given that CD44 appears to be required in a strain-transcendent manner, future studies using a diverse set of strains will also be needed to comprehensively define *P. falciparum* binding partners for CD44, which may extend to other members of the EBA/PfRH families.

EBA-175 and the other alternative ligands have well-defined roles during the initial phase of invasion, when they mediate merozoite apical reorientation and RBC membrane deformation through interactions with the glycophorins and CR1. The discovery that CD44 helps to facilitate EBA-175-induced signaling to the RBC cytoskeleton has broadened our understanding of the molecular events occurring during invasion. Our findings here show that CD44 cross-linking enhances invasion, likely through signaling to the cytoskeleton and altering the accessibility of BSG and other receptors. Together, these insights support a model in which initial engagement of CD44 by EBA proteins localized at the apical end of the invading merozoite serves to alter the RBC in a manner that enhances the interaction between the PCRCR complex and BSG and primes the cell for parasite internalization.

## ACKNOWLEDGEMENTS

The authors thank Ben Seager from the Cowman lab at WEHI, John Liao from Applied Biomics, Inc. (Hayward, CA), the International Blood Group Reference Laboratory, and Matthias Garten from Stanford University for technical assistance. The authors are grateful to Zihuai He, Paul Bollyky, members of the Egan Laboratory, John Boothroyd, Matthias Garten, and members of their respective labs for helpful discussions. This work was supported, in part, by NIH National Heart Lung and Blood Institute Grant R01HL166249 (ESE) and a Bridge Grant Award from the American Society of Hematology (ESE). AKK was funded by a T32 training grant in pediatric nonmalignant hematology and stem cell biology (DK098132-06) as well as postdoctoral support fellowship from Stanford Maternal and Child Health Research Institute. ESE is a Chan Zuckerberg Biohub San Francisco investigator and a Tashia and John Morgridge Endowed Faculty Scholar of the Stanford Maternal and Child Health Research Institute. NHT and NDS are supported by the Intramural Research Program of the NIH, National Institute of Allergy and Infectious Diseases.

## REFERENCES

1. World Malaria Report 2023. World Health Organization; 2023.

2. Cowman AF, Crabb BS. Invasion of Red Blood Cells by Malaria Parasites. Cell. 2006;124(4):755–766. doi:10.1016/j.cell.2006.02.006

3. Cowman AF, Tonkin CJ, Tham WH, Duraisingh MT. The Molecular Basis of Erythrocyte Invasion by Malaria Parasites. Cell Host & Microbe. 2017;22(2):232–245. doi:10.1016/j.chom.2017.07.003

4. Paul AS, Egan ES, Duraisingh MT. Host–parasite interactions that guide red blood cell invasion by malaria parasites: Current Opinion in Hematology. 2015;22(3):220–226. doi:10.1097/MOH.0000000000000135

5. Crosnier C, Bustamante LY, Bartholdson SJ, et al. Basigin is a receptor essential for erythrocyte invasion by Plasmodium falciparum. Nature. 2011;480(7378):534–537. doi:10.1038/nature10606

6. Farrell B, Alam N, Hart MN, et al. The PfRCR complex bridges malaria parasite and erythrocyte during invasion. Nature. 2024;625(7995):578–584. doi:10.1038/s41586-023-06856-1

7. Jamwal A, Constantin CF, Hirschi S, et al. Erythrocyte invasion-neutralising antibodies prevent Plasmodium falciparum RH5 from binding to basigin-containing membrane protein complexes. Silvie O, Soldati-Favre D, Blackman MJ, eds. eLife. 2023;12:e83681. doi:10.7554/eLife.83681

8. Scally SW, Triglia T, Evelyn C, et al. PCRCR complex is essential for invasion of human erythrocytes by Plasmodium falciparum. Nat Microbiol. 2022;7(12):2039–2053. doi:10.1038/s41564-02201261-2

9. Wong W, Huang R, Menant S, et al. Structure of Plasmodium falciparum Rh5–CyRPA–Ripr invasion complex. Nature. 2019;565(7737):118–121. doi:10.1038/s41586-018-0779-6

10. Richard D, MacRaild CA, Riglar DT, et al. Interaction between Plasmodium falciparum Apical Membrane Antigen 1 and the Rhoptry Neck Protein Complex Defines a Key Step in the Erythrocyte Invasion Process of Malaria Parasites. Journal of Biological Chemistry. 2010;285(19):14815–14822. doi:10.1074/jbc.M109.080770

11. Weiss GE, Gilson PR, Taechalertpaisarn T, et al. Revealing the Sequence and Resulting Cellular Morphology of Receptor-Ligand Interactions during Plasmodium falciparum Invasion of Erythrocytes. PLOS Pathogens. 2015;11(2):1–25. doi:10.1371/journal.ppat.1004670

12. Koch M, Baum J. The mechanics of malaria parasite invasion of the human erythrocyte – towards a reassessment of the host cell contribution. Cellular Microbiology. 2016;18(3):319–329. doi:10.1111/cmi.12557

13. Egan ES, Jiang RHY, Moechtar MA, et al. A forward genetic screen identifies erythrocyte CD55 as essential for Plasmodium falciparum invasion. Science. 2015;348(6235):711–714. doi:10.1126/science.aaa3526

14. Baro B, Kim CY, Lin C, et al. Plasmodium falciparum exploits CD44 as a coreceptor for erythrocyte invasion. Blood. 2023;142(23):2016–2028. doi:10.1182/blood.2023020831

15. Salinas N, Tang W, Tolia N. Blood-Stage Malaria Parasite Antigens: Structure, Function, and Vaccine Potential. Journal of molecular biology. Published online 2019. doi:10.1016/j.jmb.2019.05.018

16. Sim BKL, Chitnis CE, Wasniowska K, Hadley TJ, Miller LH. Receptor and Ligand Domains for Invasion of Erythrocytes by Plasmodium falciparum. Science. 1994;264(5167):1941–1944. doi:10.1126/science.8009226

17. Maier AG, Duraisingh MT, Reeder JC, et al. Plasmodium falciparum erythrocyte invasion through glycophorin C and selection for Gerbich negativity in human populations. Nat Med. 2003;9(1):87–92. doi:10.1038/nm807

18. Paing MM, Salinas N, Adams Y, et al. Shed EBA-175 mediates red blood cell clustering that enhances malaria parasite growth and enables immune evasion. eLife. 2018;7. doi:10.7554/eLife.43224

19. Koch M, Wright KE, Otto O, et al. Plasmodium falciparum erythrocyte-binding antigen 175 triggers a biophysical change in the red blood cell that facilitates invasion. Proceedings of the National Academy of Sciences. 2017;114:4225–4230. doi:10.1073/pnas.1620843114

20. Sisquella X, Nebl T, Thompson JK, et al. Plasmodium falciparum ligand binding to erythrocytes induce alterations in deformability essential for invasion. eLife. 2017;6. doi:10.7554/eLife.21083

21. Salinas N, Tolia N. A quantitative assay for binding and inhibition of Plasmodium falciparum Erythrocyte Binding Antigen 175 reveals high affinity binding depends on both DBL domains. Protein expression and purification. 2014;95:188–194. doi:10.1016/j.pep.2013.12.008

22. Alanine D, Quinkert D, Kumarasingha R, et al. Human Antibodies that Slow Erythrocyte Invasion Potentiate Malaria-Neutralizing Antibodies. Cell. 2019;178:216-228.e21. doi:10.1016/j.cell.2019.05.025

23. Chen L, Lopaticki S, Riglar DT, et al. An EGF-like Protein Forms a Complex with PfRh5 and Is Required for Invasion of Human Erythrocytes by Plasmodium falciparum. PLoS Pathogens. 2011;7. doi:10.1371/journal.ppat.1002199

24. Tonkin CJ, van Dooren GG, Spurck TP, et al. Localization of organellar proteins in Plasmodium falciparum using a novel set of transfection vectors and a new immunofluorescence fixation method. Molecular and Biochemical Parasitology. 2004;137(1):13–21. doi:10.1016/j.molbiopara.2004.05.009

25. Bolte S, Cordelières FP. A guided tour into subcellular colocalization analysis in light microscopy. Journal of Microscopy. 2006;224(3):213–232. doi:10.1111/j.1365-2818.2006.01706.x

26. Kanjee U, Grüring C, Chaand M, et al. CRISPR/Cas9 knockouts reveal genetic interaction between strain-transcendent erythrocyte determinants of Plasmodium falciparum invasion. Proceedings of the National Academy of Sciences. 2017;114(44):E9356–E9365. doi:10.1073/pnas.1711310114

27. Cywes C, Wessels MR. Group A Streptococcus tissue invasion by CD44-mediated cell signalling. Nature. 2001;414(6864):648–652. doi:10.1038/414648a

28. Kim HR, Wheeler MA, Wilson CM, et al. Hyaluronan Facilitates Invasion of Colon Carcinoma Cells in Vitro via Interaction with CD44. Cancer Research. 2004;64(13):4569–4576. doi:10.1158/00085472.CAN-04-0202

29. Duraisingh MT, Maier AG, Triglia T, Cowman AF. Erythrocyte-binding antigen 175 mediates invasion in Plasmodium falciparum utilizing sialic acid-dependent and -independent pathways. Proceedings of the National Academy of Sciences. 2003;100(8):4796–4801. doi:10.1073/pnas.0730883100

30. Gaur D, Storry JR, Reid ME, Barnwell JW, Miller LH. Plasmodium falciparum Is Able To Invade Erythrocytes through a Trypsin-Resistant Pathway Independent ofGlycophorinB. Infection and Immunity. 2003;71(12):6742–6746. doi:10.1128/iai.71.12.6742-6746.2003

31. Tham WH, Wilson DW, Lopaticki S, et al. Complement receptor 1 is the host erythrocyte receptor for Plasmodium falciparum PfRh4 invasion ligand. Proceedings of the National Academy of Sciences. 2010;107(40):17327–17332. doi:10.1073/pnas.1008151107

32. Antigen CD44 Clone BRIC 222. nhsbtdbe.blob.core.windows.net

33. Aniweh Y, Gao X, Hao P, et al. P. falciparum RH5-Basigin interaction induces changes in the cytoskeleton of the host RBC. Cellular Microbiology. 2017;19(9):e12747. doi:10.1111/cmi.12747

34. Bollyky PL, Wu RP, Falk BA, et al. ECM components guide IL-10 producing regulatory T-cell (TR1) induction from effector memory T-cell precursors. Proceedings of the National Academy of Sciences. 2011;108(19):7938–7943. doi:10.1073/pnas.1017360108

35. Fanning Á, Volkov Y, Freeley M, Kelleher D, Long A. CD44 cross-linking induces protein kinase C-regulated migration of human T lymphocytes. International Immunology. 2005;17(4):449–458. doi:10.1093/intimm/dxh225

36. Kim JH, Glant TT, Lesley J, Hyman R, Mikecz K. Adhesion of Lymphoid Cells to CD44-Specific Substrata: The Consequences of Attachment Depend on the Ligand. Experimental Cell Research. 2000;256(2):445–453. doi:10.1006/excr.2000.4852

37. Larkin J, Renukaradhya GJ, Sriram V, Du W, Gervay-Hague J, Brutkiewicz RR. CD44 Differentially Activates Mouse NK T Cells and Conventional T Cells1. The Journal of Immunology. 2006;177(1):268–279. doi:10.4049/jimmunol.177.1.268

38. Nakano K, Saito K, Mine S, Matsushita S, Tanaka Y. Engagement of CD44 up-regulates Fas Ligand expression on T cells leading to activation-induced cell death. Apoptosis. 2007;12(1):45–54. doi:10.1007/s10495-006-0488-8

39. Ruppert SM, Hawn TR, Arrigoni A, Wight TN, Bollyky PL. Tissue integrity signals communicated by high-molecular weight hyaluronan and the resolution of inflammation. Immunologic Research. 2014;58(2):186–192. doi:10.1007/s12026-014-8495-2

40. Grass GD, Tolliver LB, Bratoeva M, Toole BP. CD147, CD44, and the Epidermal Growth Factor Receptor (EGFR) Signaling Pathway Cooperate to Regulate Breast Epithelial Cell Invasiveness *. Journal of Biological Chemistry. 2013;288(36):26089–26104. doi:10.1074/jbc.M113.497685

41. Slomiany MG, Grass GD, Robertson AD, et al. Hyaluronan, CD44, and Emmprin Regulate Lactate Efflux and Membrane Localization of Monocarboxylate Transporters in Human Breast Carcinoma Cells. Cancer Research. 2009;69(4):1293–1301. doi:10.1158/0008-5472.CAN-08-2491

42. Yong JJM, Gao X, Prakash P, et al. Red blood cell signaling is functionally conserved in Plasmodium invasion. iScience. 2024;27(10):111052. doi:10.1016/j.isci.2024.111052

43. Awandare GA, Spadafora C, Moch JK, Dutta S, Haynes JD, Stoute JA. Plasmodium falciparum field isolates use complement receptor 1 (CR1) as a receptor for invasion of erythrocytes. Molecular and Biochemical Parasitology. 2011;177(1):57–60. doi:10.1016/j.molbiopara.2011.01.005

44. Pasvol G, Chasis J, Mohandas N, Anstee D, Tanner M, Merry A. Inhibition of malarial parasite invasion by monoclonal antibodies against glycophorin A correlates with reduction in red cell membrane deformability. Blood. 1989;74(5):1836–1843. doi:10.1182/blood.V74.5.1836.1836

45. Shakya B, Patel SD, Tani Y, Egan ES. Erythrocyte CD55 mediates the internalization of Plasmodium falciparum parasites. Silvie O, ed. eLife. 2021;10:e61516. doi:10.7554/eLife.61516

46. Mohandas N, Gallagher PG. Red cell membrane: past, present, and future. Blood. 2008;112(10):3939–3948. doi:10.1182/blood-2008-07-161166

47. King NR, Freire CM, Touhami J, Sitbon M, Toye AM, Satchwell TJ. Basigin mediation of Plasmodium falciparum red blood cell invasion does not require its transmembrane domain or interaction with monocarboxylate transporter 1. PLOS Pathogens. 2024;20(2):e1011989. doi:10.1371/journal.ppat.1011989

48. Day CJ, Favuzza P, Bielfeld S, et al. The essential malaria protein PfCyRPA targets glycans to invade erythrocytes. Cell Reports. 2024;43(4):114012. doi:10.1016/j.celrep.2024.114012

49. Volz JC, Yap A, Sisquella X, et al. Essential Role of the PfRh5/PfRipr/CyRPA Complex during Plasmodium falciparum Invasion of Erythrocytes. Cell Host & Microbe. 2016;20(1):60–71. doi:10.1016/j.chom.2016.06.004

50. Tolia NH, Enemark EJ, Sim BKL, Joshua-Tor L. Structural Basis for the EBA-175 Erythrocyte Invasion Pathway of the Malaria Parasite Plasmodium falciparum. Cell. 2005;122(3):485. doi:10.1016/j.cell.2005.07.020

