## Supplemental data for "CD44 cross-linking promotes *Plasmodium falciparum* invasion"

### FIGURE S1.

Gating Strategy

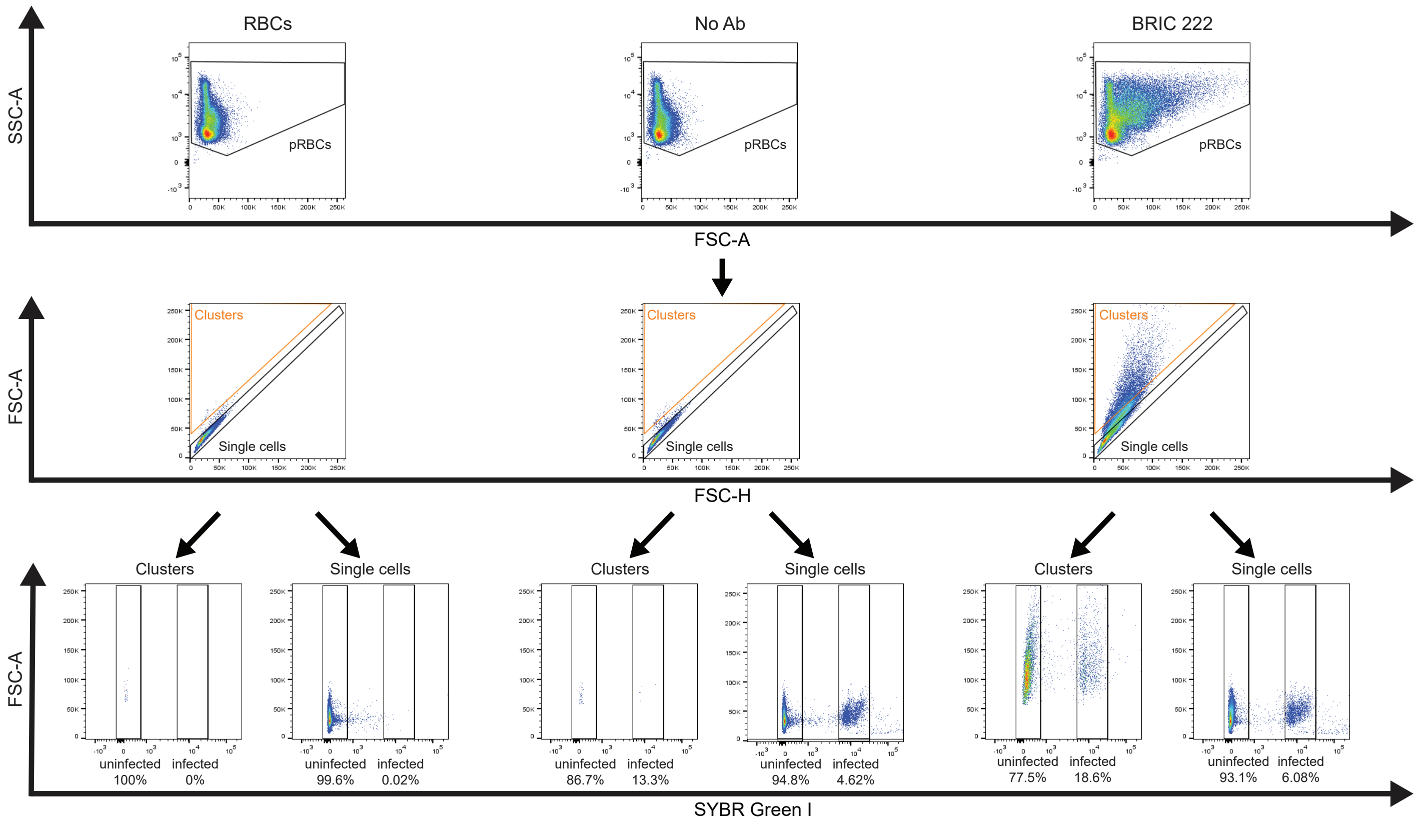

### FIGURE S2.

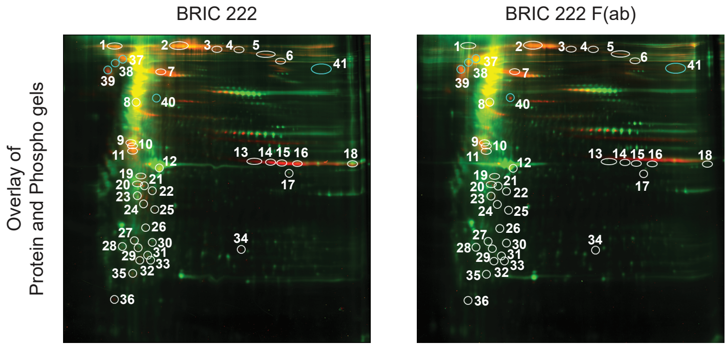


| Assigned ID | Phospho-volume BRIC 222 | Phospho-volume BRIC 222 F(ab) | Adjusted Phospho ratio (BRIC 222 F(ab)/ BRIC 222) |
|---|---|---|---|
| 1 | 1769000 | 324700 | 0.20 |
| 2 | 5980000 | 538000 | 0.10 |
| 3 | 264000 | 85700 | 0.40 |
| 4 | 271100 | 145000 | 0.65 |
| 5 | 811000 | 247000 | 0.26 |
| 6 | 1060000 | 180000 | 0.15 |
| 7 | 3810000 | 721000 | 0.19 |
| 8 | 1691000 | 147000 | 0.09 |
| 10 | 935000 | 332000 | 0.47 |
| 12 | 1100000 | 730000 | 0.65 |
| 13 | 695000 | 415000 | 0.63 |
| 14 | 1271000 | 370000 | 0.28 |
| 15 | 2043000 | 689000 | 0.32 |
| 16 | 2810000 | 1410000 | 0.47 |
| 17 | 88300 | 52700 | 0.58 |
| 18 | 895000 | 317000 | 0.33 |
| 20 | 447800 | 213800 | 0.56 |
| 21 | 297000 | 163000 | 0.67 |
| 23 | 144500 | 25500 | 0.20 |
| 24 | 235200 | 130700 | 0.62 |
| 25 | 104000 | 57500 | 0.64 |
| 27 | 79300 | 50200 | 0.64 |
| 28 | 81300 | 52400 | 0.55 |
| 30 | 56300 | 23500 | 0.38 |
| 32 | 213000 | 57100 | 0.44 |
| 35 | 227000 | 120600 | 0.66 |
| 36 | 162000 | 15900 | 0.09 |
| 37 | 446000 | 2410000 | 5.62 |
| 38 | 446000 | 1110000 | 2.33 |
| 39 | 1210000 | 2652000 | 2.28 |
| 40 | 32600 | 253000 | 8.07 |
| 41 | 44900 | 215100 | 4.79 |

### FIGURE S3.

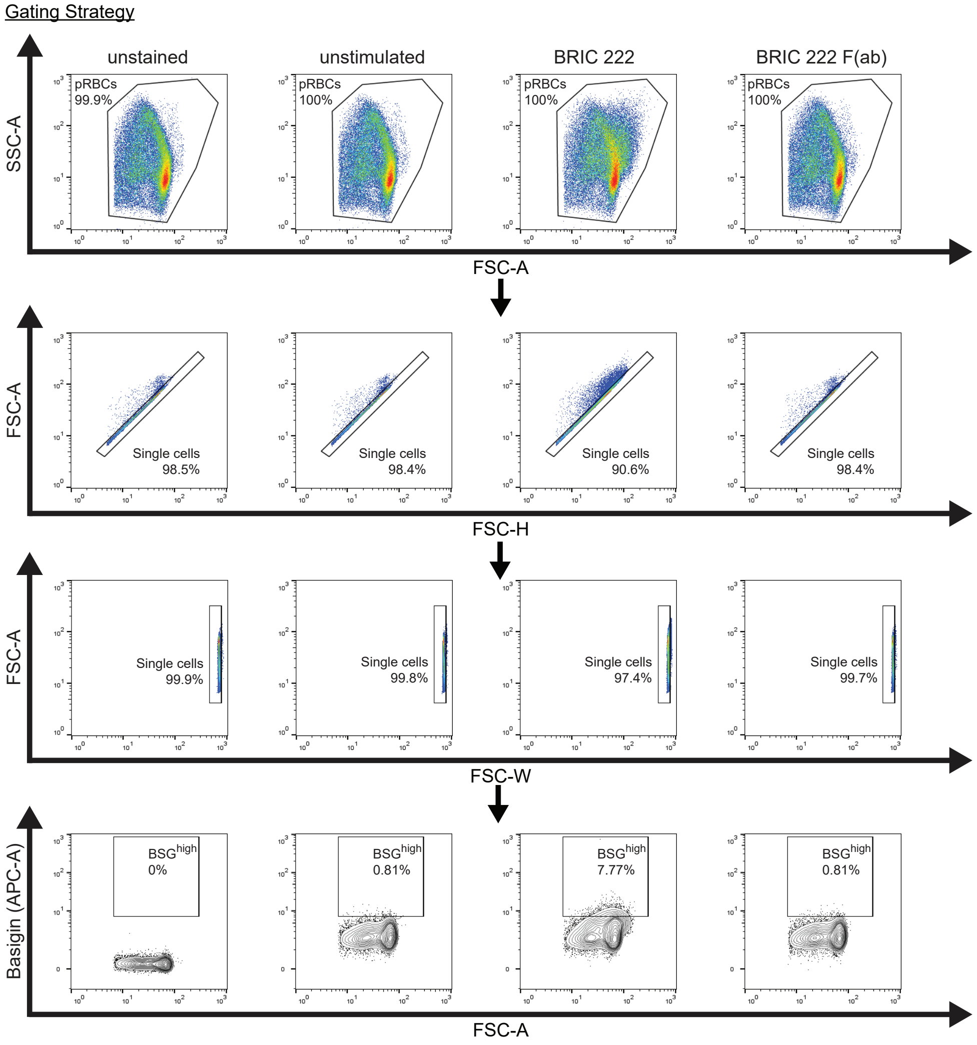

Gating Strategy
unstained
unstimulated
BRIC 222
BRIC 222 F(ab)
SSC-A
pRBCs 99.9%
pRBCs 100%
pRBCs 100%
pRBCs 100%
FSC-A
FSC-A
Single cells 98.5%
Single cells 98.4%
Single cells 90.6%
Single cells 98.4%
FSC-H
FSC-A
Single cells 99.9%
Single cells 99.8%
Single cells 97.4%
Single cells 99.7%
FSC-W
Basigin (APC-A)
BSGhigh 0%
BSGhigh 0.81%
BSGhigh 7.77%
BSGhigh 0.81%
FSC-A

### FIGURE S4.

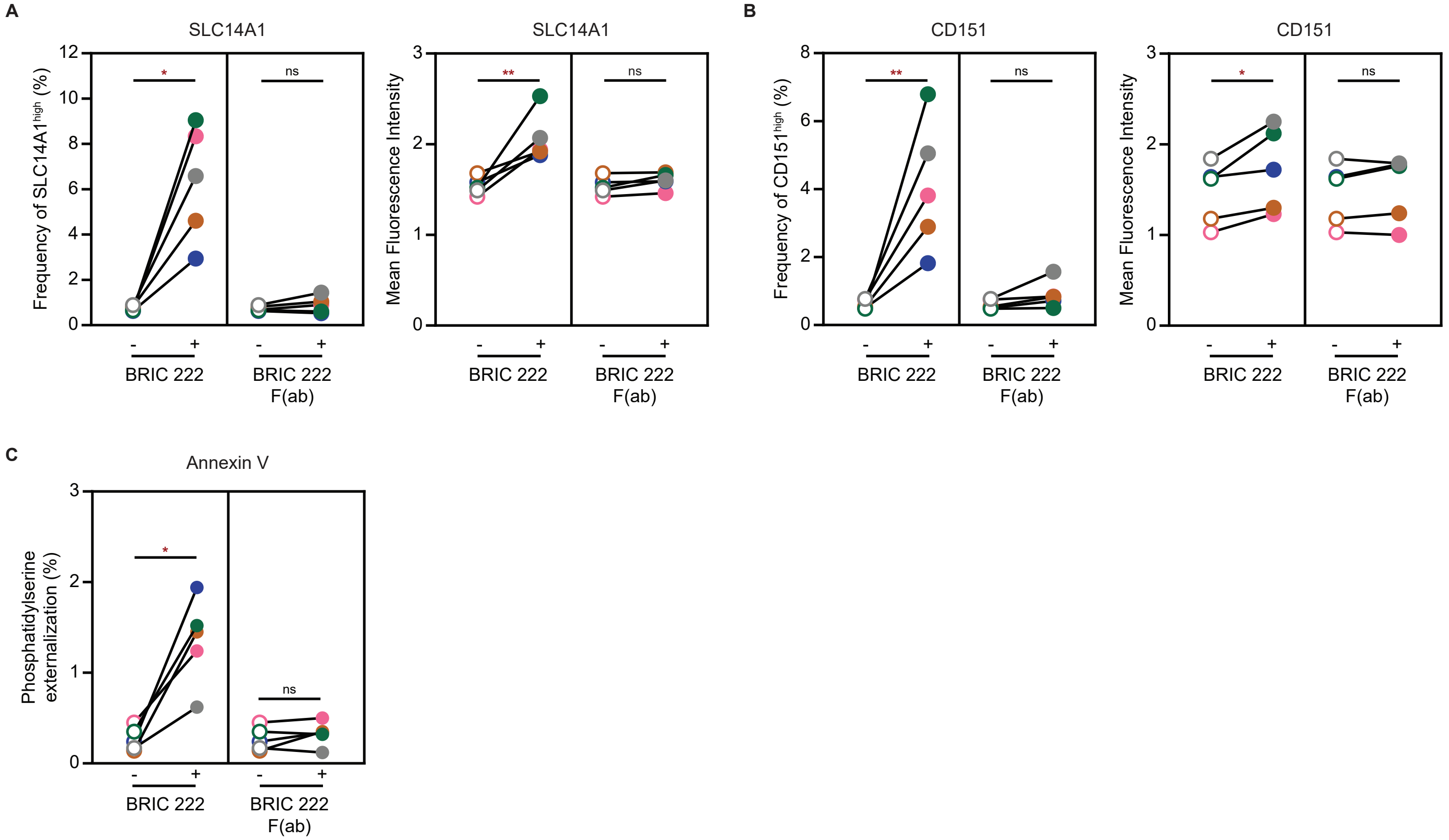
A
SLC14A1
Frequency of SLC14A1high (%)
Mean Fluorescence Intensity
BRIC 222
BRIC 222 F(ab)
ns
B
CD151
Frequency of CD151high (%)
C
Annexin V
Phosphatidylserine externalization (%)

### FIGURE S5.

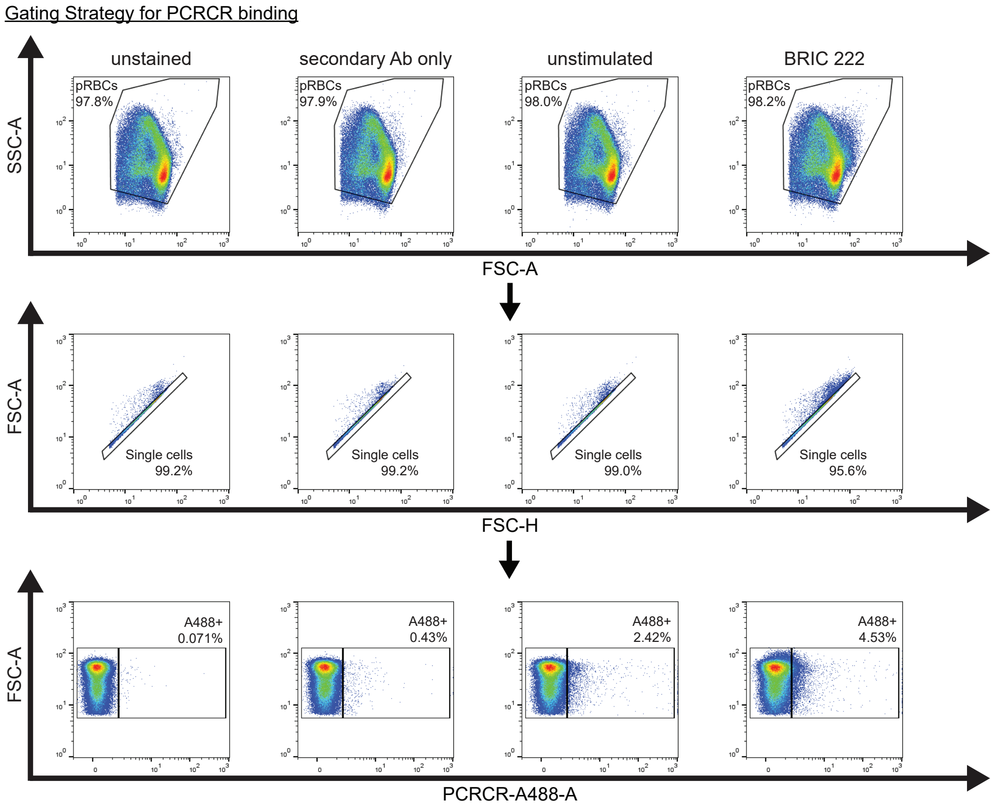

Gating Strategy for PCRCR binding
unstained
secondary Ab only
unstimulated
BRIC 222
SSC-A
pRBCs
97.8%
pRBCs
97.9%
pRBCs
98.0%
pRBCs
98.2%
FSC-A
FSC-A
Single cells
99.2%
Single cells
99.2%
Single cells
99.0%
Single cells
95.6%
FSC-H
FSC-A
A488+
0.071%
A488+
0.43%
A488+
2.42%
A488+
4.53%
PCRCR-A488-A

### FIGURE S6.

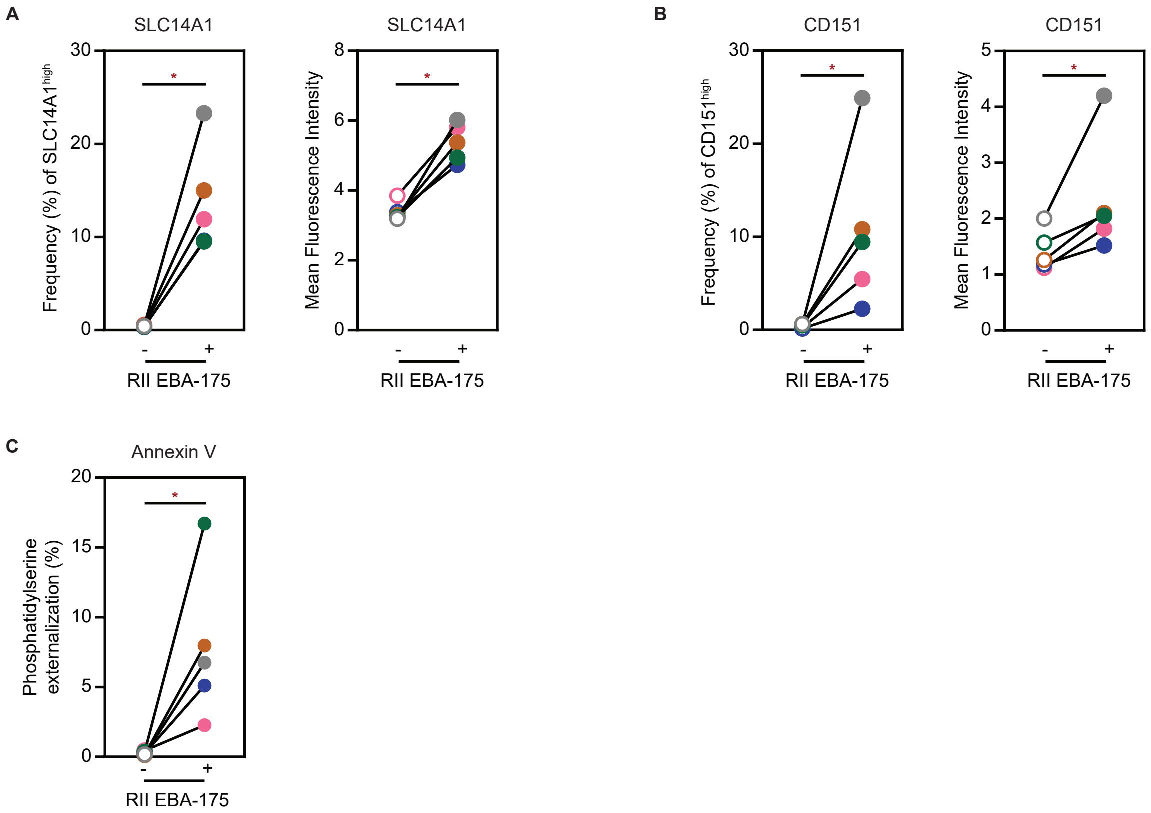
A
SLC14A1
Frequency (%) of SLC14A1^high
Mean Fluorescence Intensity
RII EBA-175
B
CD151
Frequency (%) of CD151^high
Mean Fluorescence Intensity
RII EBA-175
C
Annexin V
Phosphatidylserine externalization (%)
RII EBA-175

### FIGURE S7.

**A** Gating Strategy for PCRCR binding

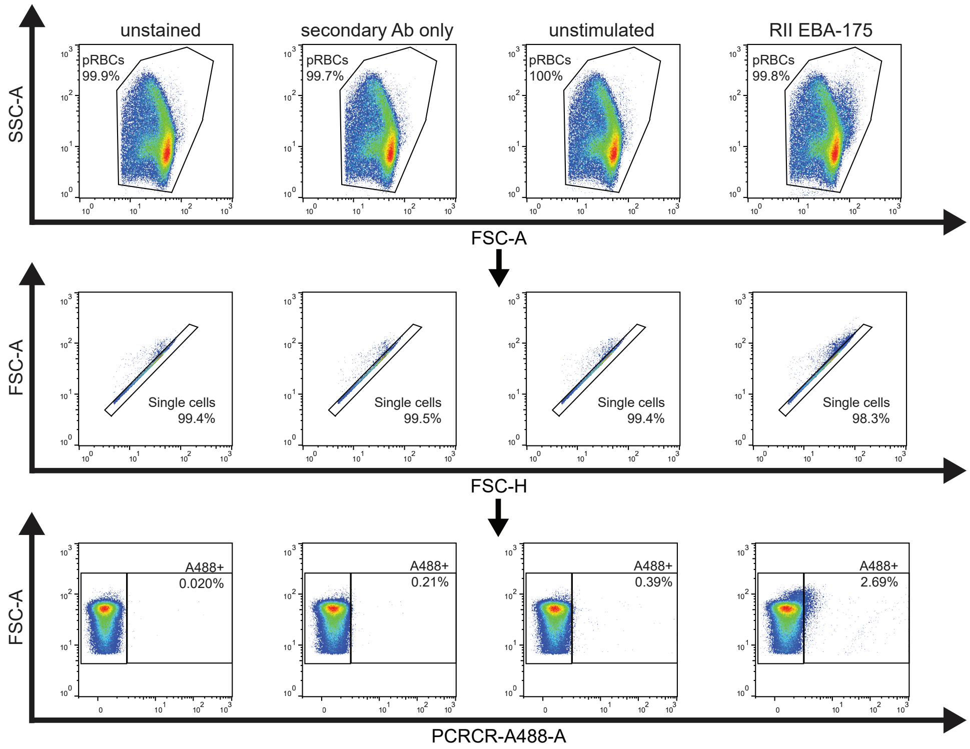


**B** Gating Strategy for PfRH5 binding

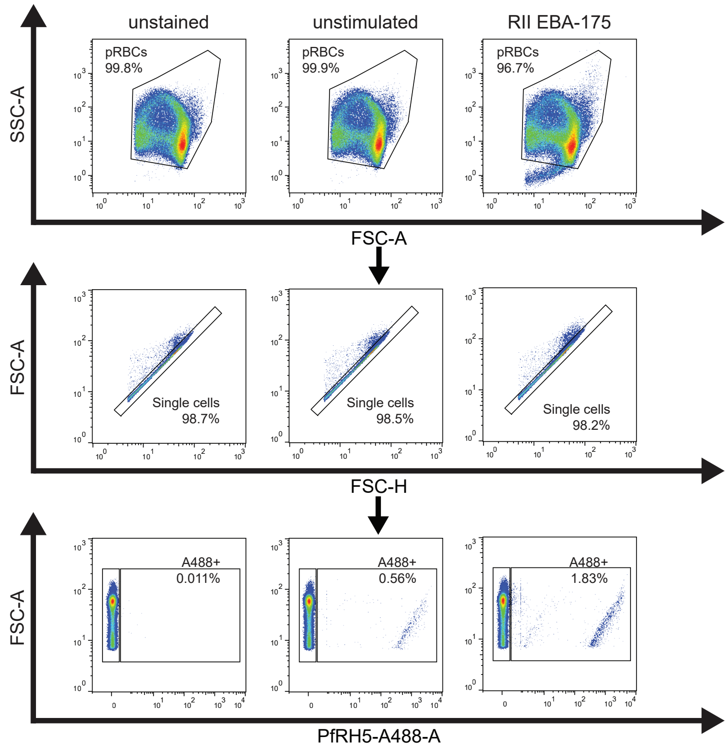

#### SUPPLEMENTAL FIGURE LEGENDS

**Figure S1: Gating strategy for invasion assays**
Gating strategy for flow cytometry analysis of *P. falciparum* invasion assays with BRIC 222 stimulation. Following RBC gating, the cells were separately gated as "Clusters" or "Single Cells" based on the forward scatter (size). Parasite infection of each cell population was then assessed by SYBR Green I nucleic acid staining. A representative flow plot is shown.

**Figure S2: CD44 cross-linking affects phosphorylation of RBC proteins**
RBCs were stimulated with BRIC 222 or BRIC 222 F(ab) at 25 μg/ml. Ghost RBCs of these stimulated samples were then generated and analyzed for their phosphorylation status by 2D-DIGE. The overlay of Total Protein and Phosphoprotein gels are shown. Top: All spots with differential phosphorylation, with ≥1.5 or ≤0.67 fold-change in phosphorylation, between the BRIC 222 F(ab)- and BRIC 222-stimulated conditions, are circled. White circles: higher phosphorylation in BRIC 222-stimulated samples. Blue circles: higher phosphorylation in BRIC 222 F(ab)-stimulated samples. Bottom: Each differentially phosphorylated spot is listed with its Phospho-volume as well as the Adjusted Phospho ratio (adjusted value based on the protein expression).

**Figure S3: Gating Strategy for BSG$^{high}$**
Gating strategy for flow cytometry analysis of Basigin (BSG) detection by the anti-BSG monoclonal antibody targeting an invasion epitope (TRA-1-85). A representative donor is shown. RBCs were stimulated with BRIC 222 or BRIC 222 F(ab) prior to surface protein detection. The samples were first gated for RBCs based on the forward and side scatter, followed by single cell gating based on forward scatter. The surface expression of BSG was then assessed. The BSG$^{high}$ population, relatively to that of unstimulated condition, is boxed with frequencies indicated. The same gating strategy was applied to all other surface markers assessed.

**Figure S4: CD44 cross-linking alters RBC membrane**
RBCs were stimulated with BRIC 222 or BRIC 222 F(ab) at 25 μg/ml and then assessed for surface protein changes, as in Figure 4. **(A-B)** Frequencies of SLC14A1$^{high}$ (A) and CD151$^{high}$ (B) and the respective MFI of five donors are plotted. **(C)** Phosphatidylserine (PS) externalization of RBC membrane following stimulation of BRIC 222 or BRIC 222 F(ab) (25 μg/ml) was measured with Annexin V. Frequency of cells with PS externalization of five donors are shown. For each graph, a color dot represents an individual donor. Statistical analysis: Friedman test with FDR correction;* $p \leq 0.05$, ** $p \leq 0.01$, ns: non-significant.

**Figure S5: Gating Strategy for PCRCR binding of BRIC 222-stimulated RBCs**
Gating strategy for flow cytometry analysis of PCRCR binding to RBCs from one representative donor is shown. The samples were first gated for RBCs based on the forward and side scatter, followed by single cell gating based on forward scatter. The PCRCR binding was detected with primary antibody against RH5 (R5.011), followed by an A488-conjugated secondary antibody. The PCRCR binding is boxed with A488+ frequencies indicated.

**Figure S6: *P. falciparum* EBA-175 alters RBC membrane**
RBCs were stimulated with 2 μM RII EBA-175 and assessed for membrane changes. Gating strategy was performed as described in Figure S3. **(A-B)** Frequencies of SLC14A1$^{high}$ (A), CD151$^{high}$ (B) and the respective MFI of five donors are plotted. **(C)** Phosphatidylserine (PS) externalization of RBC membrane was detected with Annexin V. Frequency of cells with PS externalization of five donors are shown. For

each graph, a color dot represents an individual donor. Statistical analysis: Wilcoxon matched-pairs signed rank test, one-tailed; * $p \leq 0.05$.

**Figure S7: Gating Strategy for PCRCR and PfRH5 binding of EBA-175-stimulated RBCs**
**(A-B)** Gating strategy for flow cytometry analysis of PCRCR (A) or PfRH5 (B) binding to RBCs from one representative donor each is shown. The samples were first gated for RBCs based on the forward and side scatter, followed by single cell gating based on forward scatter. The PCRCR binding was detected with primary antibody against RH5 (R5.011), followed by an A488-conjugated secondary antibody. The PfRH5 binding was detected with anti-PfRH5 (5A9-488). The percentage of PCRCR or PfRH5 binding is boxed, with the A488+ frequencies indicated.
